# Single Cell Profiling of Acute Kidney Injury Reveals Novel Transcriptional Signatures, Mixed Identities and Epithelial-to-Stromal Crosstalk

**DOI:** 10.1101/2019.12.30.890905

**Authors:** Valeria Rudman-Melnick, Mike Adam, Andrew Potter, Saagar M. Chokshi, Qing Ma, Keri A. Drake, Meredith P. Schuh, J. Matthew Kofron, Prasad Devarajan, S. Steven Potter

## Abstract

Acute kidney injury (AKI) is a rapid decline of renal function, with an incidence of up to 67% of intensive care unit patients. Current treatments are merely supportive, emphasizing the need for deeper understanding that could lead to improved therapies. We used single cell RNA sequencing, *in situ* hybridization and protein expression analyses to create comprehensive renal cell specific transcriptional profiles of multiple AKI stages. We revealed that AKI induces marked dedifferentiation, renal developmental gene activation and mixed identities in injured renal tubules. Moreover, we identified potential pathologic crosstalk between epithelial and stromal cells, and several novel genes involved in AKI. We also demonstrated the definitive effects of age on AKI outcome, and showed that renal developmental genes hold a potential as novel AKI markers. Moreover, our study provides the resource power which will aid in unraveling the molecular genetics of AKI.

## Introduction

Acute kidney injury (AKI) is a group of syndromes characterized by abrupt renal function decline associated with pronounced mortality and co-morbidities in other organs (Hoste et al., 2018). AKI is a common condition with estimated prevalence ranging from <1% to 66% and increasing trends of hospitalization number (Pavkov et al., 2018), which represents a global burden to the healthcare system. Moreover, AKI episodes can initiate or exacerbate chronic kidney disease (CKD), a debilitating condition requiring lifelong dialysis or renal transplantation (Hsu and Hsu, 2016). Thus, examining the molecular signaling pathways involved in early AKI response is paramount for treatment development.

Single cell RNA sequencing (scRNA-seq) has been used to thoroughly characterize the gene expression profiles of normal cells types of the developing and adult kidney for both mouse and human (Brunskill et al., 2014; Combes et al., 2019; Hochane et al., 2019; Ransick et al., 2019; Wang et al., 2018). In this study, we present the first comprehensive atlas of renal expression changes at multiple stages of AKI progression at the single cell level. Since disrupted renal perfusion represents one of the most clinically relevant causes of kidney disease (Yang et al., 2010), we used the unilateral ischemia/reperfusion (UIR) model of AKI which allows for inducing severe renal injury without significant mortality (Le Clef et al., 2016). We identified unique injury response gene expression signatures of all renal cell populations. Proximal tubule cells showed a rapid reduced expression of differentiation markers such as *Slc34a1*, while showing increased expression of genes normally expressed during development, such as *Sox4* and *Cd24a*. Particularly striking, many of the injured cells showed simultaneous expression of markers of multiple nephron cell types, consistent with reversion to an early developmental stage (Brunskill et al., 2014; Magella et al., 2018). We identified several novel genes which were not previously reported to play a role in AKI response. Further, we found that injured tubules expressed genes normally associated with the stromal lineage. Analysis of ligand/receptor interactions revealed potential pathologic crosstalk between epithelial and stromal cells. We also examined the AKI response as a function of age of injury, finding that young mice show a much more rapid recovery with greatly reduced long term fibrosis. The results show that renal developmental genes represent potential novel AKI markers.

## Results and Discussion

### Single Cell RNA Sequencing Reveals Proximal Tubule Dedifferentiation and Mixed Identity Cells in the Ischemia Reperfusion induced AKI

To examine the molecular and cellular nature of acute kidney injury (AKI) response, we used the clinically relevant model of Unilateral Ischemia/Reperfusion (UIR) AKI (Le Clef et al., 2016) (Figure 1A). To evaluate the transcriptional changes in renal cell populations over the course of AKI, we performed single cell RNA-seq (scRNA-seq) on renal cell suspensions from Swiss-Webster 4 weeks old male mice at Days 1, 2, 4, 7, 11 and 14 after UIR (Figure 1B). We carried out scRNA-seq using Drop-Seq as previously described (Adam et al., 2017; Macosko et al., 2015; Magella et al., 2018), and used unsupervised clustering (Stuart et al., 2019) to generate Uniform Manifold Approximation and Projection (UMAP) plots which separated the cells into distinct groups (Fig. 1C). We identified resulting cell clusters including *Nphs2*-positive podocytes, proximal tubule (*Slc34a1*), distal tubule (*Slc12a3*), loop of Henle (*Slc12a1*), collecting duct (*Aqp2* for principal and *Foxi1* for intercalated cells) as well as endothelial cells (*Pecam1*), pericytes (*Pdgfrβ*), macrophages (*Cd68*), T cells (*Cd3g*), and stromal/pericytes cells (*Pdgfrb*) (Figure 1C and S1A).

**Figure 1.**
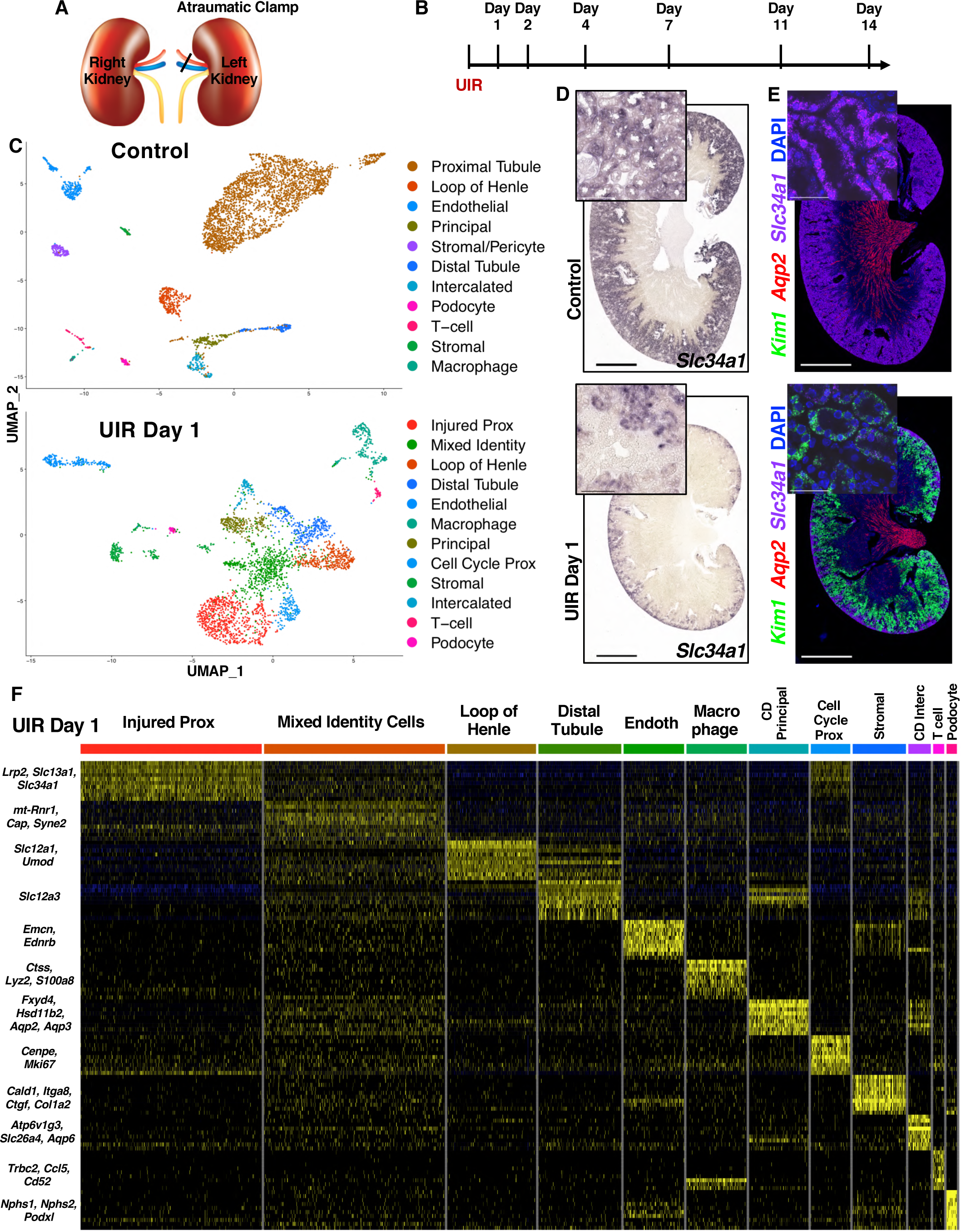
Single Cell RNA Sequencing Reveals Proximal Tubule Dedifferentiation and Mixed Cell Identity in the Ischemia Reperfusion induced AKI. (A) UIR procedure scheme. (B) The experimental timeline. (C) UMAP plots show the renal cell populations in the Control and UIR Day 1. (D) *Slc34a1* CISH, Control and UIR Day 1, 4x, 40x. (E) *Kim1* (green), *Aqp2* (red), *Slc34a1* (purple) RNAscope, Control and UIR Day 1, 4x, 60x. (F) Heatmap shows the relative marker gene expression in UIR Day 1 of renal cell types. Endoth, Endothelial; CD, Collecting Duct; Interc, Intercalated. The Mixed Identity Cells show stochastic expression of markers of many different cell types. Scale: 4x – 2500*µ*m, 40x – 100*µ*m, 60x – 25*µ*m. Related to Figures S1, S2, Table S1.

We first compared gene expression patterns of control and UIR Day 1 kidneys for each cell type (Table S1). In the normal kidney, the renal sodium-dependent phosphate transporter *Slc34a1* marks mature proximal tubules, which represent the predominant cell type in the renal cortex, as shown by both scRNA-seq and chromogenic *in situ* hybridization (CISH) (Figures 1C and 1D, S1A). UIR, however, resulted in a markedly reduced *Slc34a1-*positive mature proximal tubule population (Figures 1C and S1B). We validated this finding with CISH performed on an independent cohort of 4 weeks old Swiss-Webster mice treated with the identical UIR. While the control mice abundantly expressed *Slc34a1* in the cortex and inner medulla, the UIR Day 1 exhibited significantly reduced *Slc34a1* levels, with remaining expression only in the outer cortex (Figure 1D). This observation is consistent with the previously reported injury-induced proximal tubule dedifferentiation (Kramann et al., 2015; Kusaba et al., 2014). Moreover, we identified the abundant elevation of *kidney injury molecule-1* (*Kim1*, a.k.a. *Havcr1*) in the cortex and outer medulla of the UIR Day 1 kidney compared to control, using fluorescent *in situ* hybridization (FISH, a.k.a. RNAscope) (Figure 1E). Kim1 is a clinically recognized AKI marker (Khreba et al., 2019); as we observed and others (Bonventre, 2008; Gauer et al., 2016) have shown, *Kim1* is predominantly expressed in the UIR Day 1 injured proximal tubules and is nearly absent in the Control (Figure S1A). Based on *Kim1* and *Slc34a1* distribution, we identified the cluster of “Injured Prox”, defined by the most prominent *Kim1* and lowered *Slc34a1* levels (Figure 1C, S1B). The cluster of “Cell Cycle Prox” was identified based on the high expression of cell cycle markers, including the proliferation marker *Mki67* (Figure S1B). Next, we identified the loop of Henle, distal tubule, collecting duct and found that they all are highly enriched with *Lipocalin 2* (*Lcn2*), another clinically used AKI marker specific for the distal segments of the nephron tubule (Schmidt-Ott et al., 2007; Singer et al., 2016; Sun et al., 2018) (Figures S1B and S1C). Histological comparison of UIR Day 1 vs Control using Hematoxylin & Eosin (H&E) staining revealed signs of pronounced tubular epithelial injury, including renal tubular dilation, epithelial flattening and cast formation (Figure S1D). Thus, we showed that UIR Day 1 comprises all classical AKI features, including marked mature cell dedifferentiation and injury in multiple nephron tubule segments (Basile et al., 2012; Kudose et al., 2018).

To identify the molecular signaling pathways and biological processes which might play a role in the early AKI response, we performed functional enrichment gene analysis and candidate gene prioritization using the ToppGene Suite (Chen et al., 2009). Gene Ontology (GO) analysis of UIR Day 1 injured proximal tubules showed substantial reduction of proximal tubule metabolism, including oxidation-reduction processes (*Slc37a4, Slc25a2, Aco1, Aco2, Cyp2e1, Miox*) and fatty acid catabolism (*Acadl, Acadm, Cd36, Cpt1a, Crot, Slc27a2*), which are the key proximal tubule metabolic pathways (Kang et al., 2015; Simon and Hertig, 2015) (Figure S1E, Table S1). We also observed the UIR induced reduction of transmembrane transport pathways (*Slc34a1, Aqp1, Lrp2, Slc7a13, Slc22a6*) crucial for normal proximal tubule homeostatic functions. Conversely, UIR caused elevation of apoptotic processes (*Acin1, Tnfrsf12a, Clu, Nfkbia, Lgals1, Ctsd*), cell cycle regulation (*Ccnd1, Ccnl1, Ctnnb1, Pcna*), granulocyte activation and cytokine response (*C3, S100a11, Lcn2, Cstb, Anxa2)*, and cellular response to oxygen (*Mgst1, Gpx3, Aoc1*) in the “Injured Prox” (Figure S1E, Table S1).

Notably, we found that AKI not only induces dedifferentiation, but causes an ambiguous transcriptional identity and ectopic expression of mature defined renal cell populations markers. In the UIR Day 1, we noticed a large population of cells with overlapping expression of the collecting duct marker *Aquaporin 2* (*Aqp2)* (Yu et al., 2014) and the loop of Henle marker *Uromodulin* (*Umod)* (Tokonami et al., 2018) (Figure S2A). These “Mixed Identity Cells” were located between proximal tubules, loop of Henle, distal tubule and collecting duct on the UMAP (Figures 1C, S2A), suggesting the intermediate character. Also, the scRNA-seq data showed that many of these cells expressed both *Kim1* and *Lcn2* (Figure S2B) which mark proximal and distal nephron tubule segment injury, respectively. A heat map revealed the apparent mixed identity, showing that these cells exhibit stochastic expression of markers of many different kidney cell types (Figure 1F), including principal (*Fxyd4, Hsd11b2, Aqp2, Aqp3*) and intercalated (*Atp6v1g3, Atp6v0d2, Slc26a4, Aqp6*) cells of the collecting duct (Chen et al., 2017), proximal tubule (*Slc34a1*), loop of Henle (*Slc12a1, Umod*) and distal tubule (*Slc12a3*), which was not observed in the control (Figure S2C). This scRNA-seq observation was reproduced in an additional independent cohort of validation animals. To detect the mixed identity cells, we hybridized *Slc34a1, Slc12a1, Slc12a3* and *Aqp2* RNAscope probes to the UIR Day 1 and Control kidney sections and found that UIR result in the remarkable presence of cells expressing multiple epithelial compartments markers. Prominently, IMARIS quantification revealed significantly increased numbers of *Aqp2, Slc12a3* and *Slc12a1* transcripts in the tubules exhibiting lowered *Slc34a1* levels (Figures 2A–2D), as predicted by scRNA-seq. The presence of *Aqp2* transcripts in *Slc34a1*-positive tubules demonstrates the marked plasticity of adult renal injured cells, since the proximal tubule (*Slc34a1*) and the collecting duct (*Aqp2*) arise from the different developmental lineages, the metanephric mesenchyme and the nephric duct respectively (Costantini and Kopan, 2010; McMahon, 2016). Of interest, we also noticed that AKI not only induces the presence of the mixed identity cluster, but also results in inappropriate ectopic gene expression in cells that clustered as a defined cell type. For example, the RNAscope showed elevated *Aqp2* transcripts in the *Kim1*-positive injured proximal tubules (Figure 2E).

**Figure 2.**
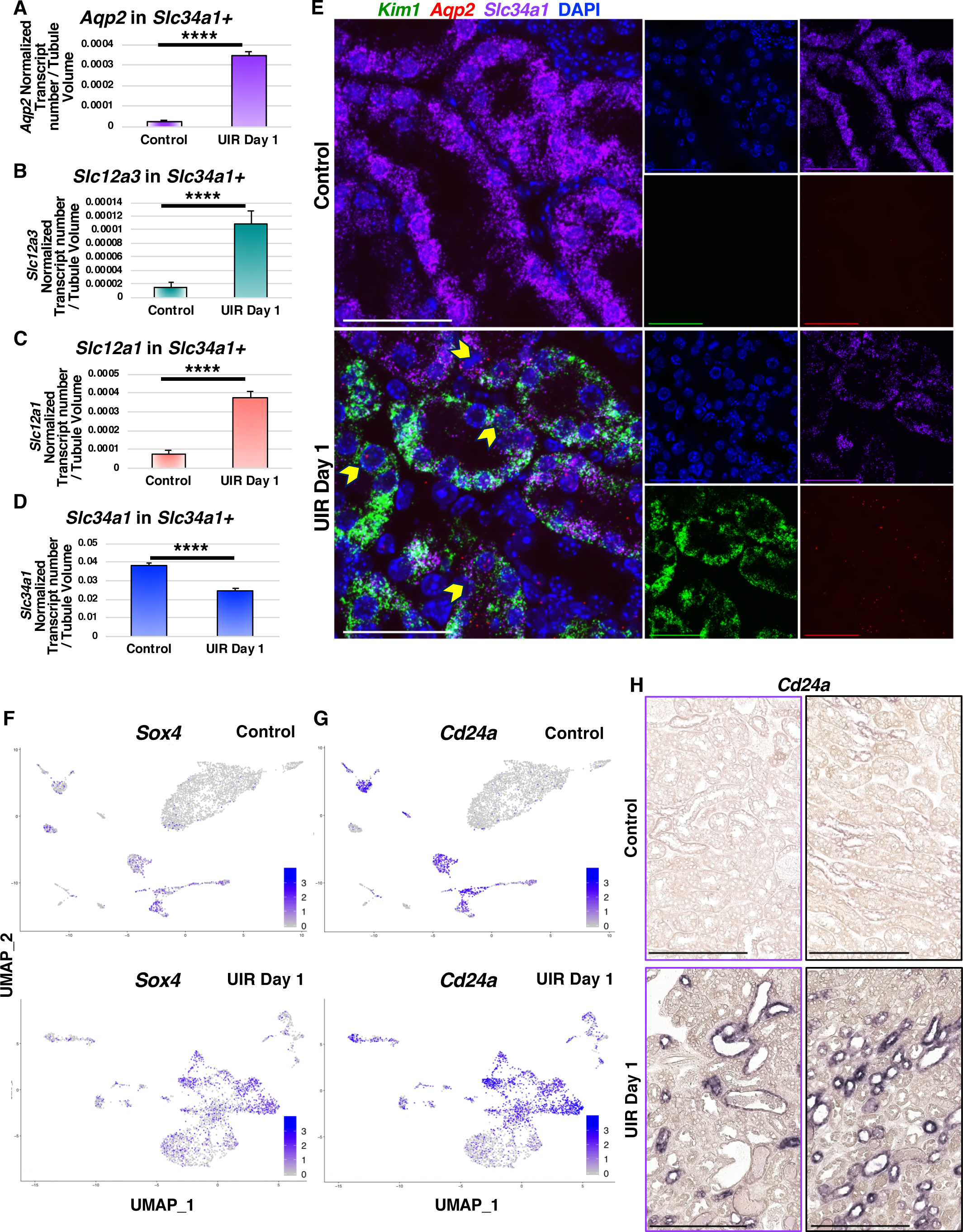
AKI Induces Renal Developmental Program Re-Activation in the Adult Kidney. (A-D) IMARIS quantification of *Aqp2, Slc12a3, Slc12a1 and Slc34a1* in the UIR Day 1 vs Control renal tubules, n=12 Z-stacks (50-70 tubules) per group, Mean ± SEM, Student’s *t* test, **** p<0.0001. (E) Representative images of *Kim1* (green), *Aqp2* (red), *Slc34a1* (purple) RNAscope, *Aqp2* in *Slc34a1-*or *Kim1-*positive UIR Day 1 proximal tubules (yellow pointers). Maximal Intensity Projection from Z-stacks, 60x 1.5 Nyquist zoom, scale 50*µ*m. (F and G) UMAPs show the elevated *Cd24a* and *Sox4* expression in UIR Day 1. Cell cluster identities are as defined in Figure 1. (H) CISH shows the elevated *Cd24a* expression in the UIR Day 1 renal tubules; 40x, cortex, purple frames (left); medulla, black frames (right); scale 100*µ*m. Related to Figure S2, Table S1.

Noteworthy, the AKI also resulted in ambiguous epithelial/stromal identities. The heat map shows that the “Mixed Identity Cells” exhibit increased expression of stromal genes, particularly activated fibroblast markers (*Ctgf, Col1a2, Cald1*) (Shieh et al., 2019; Wu et al., 2018; Zhang et al., 2017b) (Figure 1F) and genes related to the cell shape change and extracellular matrix remodeling (*Sparc, Lgals3, Spp1, Myh9, S100a10*) (Table S1), giving evidence for a more mesenchymal phenotype acquired by these cells in response to the injury. Epithelial-to-mesenchymal plasticity is a crucial contributor to the kidney injury and fibrosis (Humphreys, 2018; Lovisa et al., 2016). In contrast, this cluster exhibited much weaker expression of genes associated with more distantly related cell types, like macrophages and T-cells (Figure 1F). Overall, our data reveals that AKI induces apparent cellular plasticity between adult renal cell types. We previously showed that lineage infidelity, or mixed cell identity, occurs in the developing kidney, with the unexpected stochastic expression of markers of multiple differentiated renal cell types in single cells of the E11.5 MM, cap mesenchyme and later renal vesicles (Brunskill et al., 2014; Magella et al., 2018). However, this cellular plasticity between renal epithelial compartments, including those of different developmental origins, to our knowledge had not been previously observed in the adult AKI.

### AKI induces Renal Developmental Program Re-Activation in the Adult Kidney

Next, we further analyzed the gene expression signatures of injured renal cells. The GO analysis of the “Mixed Identity Cells” revealed the enrichment of renal developmental pathways (Figure S2C, Table S1). Notably, we identified the *Transcription Factor SRY-related HMG Box-4* (*Sox4*) (Figure 2F), implicated in the urogenital system, kidney and epithelium development by GO analysis. Previous work showed that *Sox4* is strongly expressed in nephron progenitors and ureteric bud during early and late renal development as well as in the newborn kidney (Huang et al., 2013; Yu et al., 2012). Moreover, *Six2Cre;Sox4* mutants exhibit reduced nephron number and proteinuria postnatally, which progresses to ESRD defined by glomerulosclerosis and proximal tubular injury (Huang et al., 2013), showing that *Sox4* is essential for proper nephrogenesis. However, *Sox4* has no known role in AKI. We also observed that that the “Mixed Identity Cells” were enriched with *Signal Transducer Cd24* (*Cd24a*) (Figure 2G, Table S1), which encodes a cell-surface sialoglycoprotein expressed in the uninduced metanephric mesenchyme (MM) of the developing kidney (Challen et al., 2004). Human Cd24 is also implicated in kidney development and tubular epithelial differentiation (Ivanova et al., 2010; Lazzeri et al., 2007). We observed that *Cd24a* is also elevated in the adult injured proximal tubules (Figure 2G), consistent with previous reports (Kramann et al., 2015; Kusaba et al., 2014). Moreover, the UMAPs showed that both *Sox4* and *Cd24a* were also enriched in the loop of Henle and distal tubule, and in the principal and intercalated cells of the collecting duct (Figures 2F and 2G). With CISH, we validated the elevation of *Cd24a* in both cortical and medullary renal tubules (Figure 2H). Noticeably, the AKI induced *Sox4* and *Cd24a* elevation recapitulated their developing kidney spatial expression patterns (Harding et al., 2011; McMahon et al., 2008; Yu et al., 2012), which demonstrates the profound nature of renal developmental program re-activation and mature renal cell dedifferentiation caused by the injury.

Next, we examined cell populations and molecular changes which occur later after AKI induction. Analysis of cell clusters from UIR Day 2 revealed persistent kidney injury, indicated by continued elevated *Kim1* expression and the decreased number of strongly positive *Slc34a1* proximal tubules (Figures 3A–3C). scRNA-seq also revealed the Mixed Identity cluster, characterized by marked gene expression heterogeneity and renal developmental gene elevation, along with the clusters of macrophages and T-cells, showing the enduring inflammation (Figure 3A). Histological analysis showed exacerbated renal tubular damage, compared to day 1, exemplified by pronounced epithelial flattening, tubular casts and dilation (Figure S3A). Renal tubular injury persisted through UIR Day 4 and 7; however, we observed signs of recovery, including increased *Slc34a1* expression and reduced total kidney Kim1 and Lcn2 RNA and protein levels (Figures 3A–3C, S3A-S3C). Consistent with previous work showing the role of macrophages in kidney injury resolution (Lee et al., 2011), we observed that both timepoints exhibited large macrophage infiltration (Figure 3A). The mixed identity cells were no longer present at Day 7; however, we identified the “Dedifferentiated Prox” cluster (Figure 3A, Table S2), which expressed the proximal tubular genes *Aqp1*, a water-transporting protein (Hara-Chikuma and Verkman, 2006), *Slc7a12*, a cationic amino acid transporter (Rinn et al., 2004), and *Cdh2 (N-Cadherin)* (Nurnberger et al., 2010; Prozialeck et al., 2004). However, this cluster also showed a strong renal developmental signature, including the elevation of *Sprouty1, Nephronectin*, and the *Forkhead-box transcription factor Foxc1*, which all are crucial for the kidney induction (Basson et al., 2005; Kume et al., 2000; Linton et al., 2007). scRNA-seq also showed increased *Notch2*, a crucial regulator of nephron induction, segmentation and proximal tubule epithelium development (Chung et al., 2016; Sirin and Susztak, 2012), and Notch ligand *Jag1*, in the “Dedifferentiated Prox” (Table S2). Notably, recent work from the Susztak group shows the role of Notch2 in the collecting duct cell plasticity associated with metabolic acidosis (Park et al., 2018). This population also expressed *Osr2*, which is present in the early proximal tubule, but not in the adult cortical tubule (Harding et al., 2011; McMahon et al., 2008); and the metanephric mesenchyme marker *Cd24a* mentioned above. This developmental signature gives evidence for a dedifferentiated proximal tubule state.

**Figure 3.**
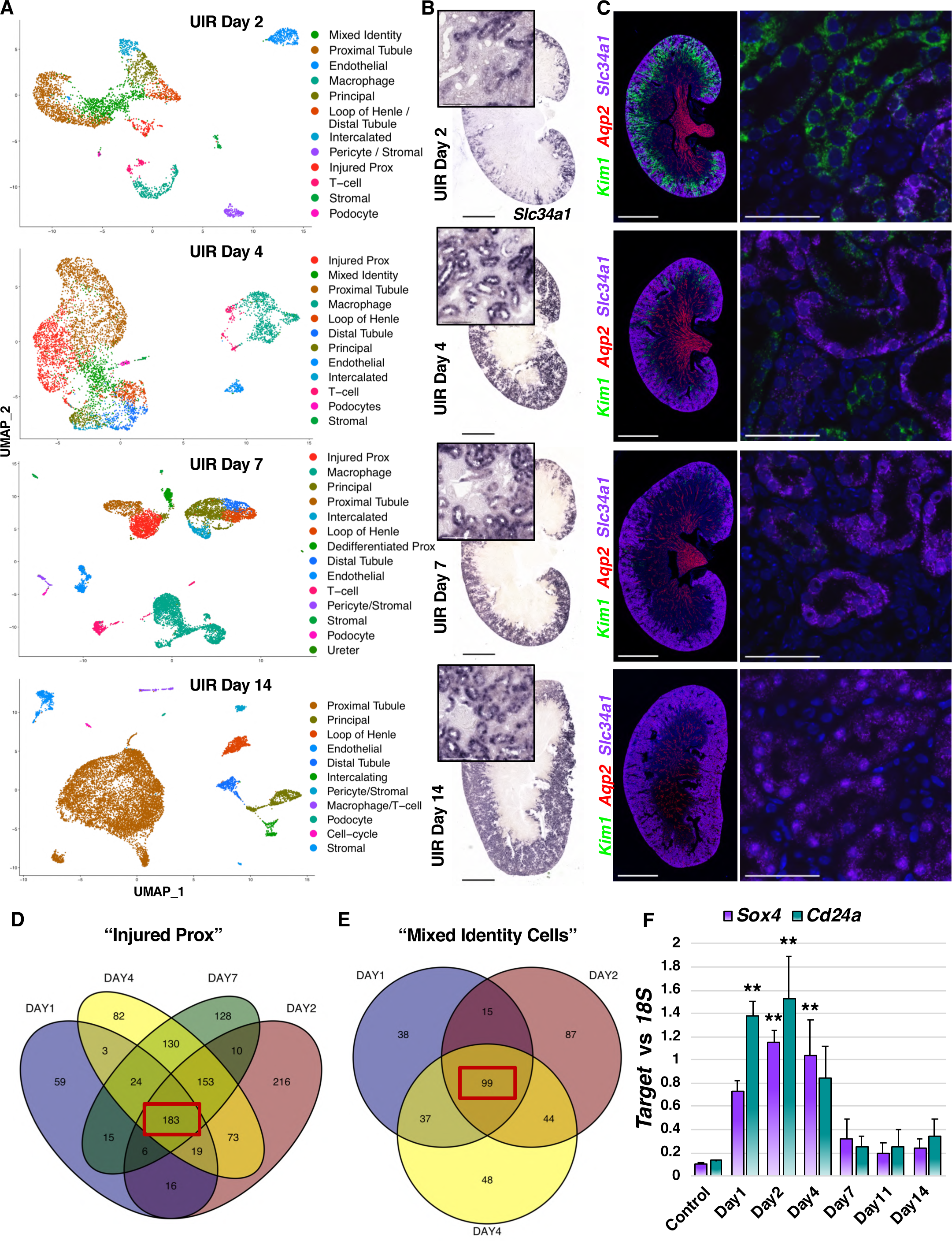
scRNA-seq Reveals a Persistent Developmental Signature over the Course of AKI. (A) UMAPs show renal cell populations in the UIR Day 2, 4, 7 and 14. Day 11 is not shown, as it was very similar to Day 14. (B) *Slc34a1* CISH, UIR Day 2, 4, 7 and 14, 4x, 40x. (C) *Kim1* (green), *Aqp2* (red), *Slc34a1* (purple) RNAscope, UIR Day 2, 4, 7 and 14. 4x, 60x. (D) Venn diagram shows the genes elevated in the “Injured Prox” at UIR Day 1, 2, 4 and 7. Injured proximal tubule gene expression patterns were compared to control proximal tubule. (E) Venn diagram shows the genes elevated in the “Mixed Identity Cells” at UIR Day 1, 2 and 4. Mixed identity cell gene expression patterns were compared to control proximal tubule. (F) qPCR shows *Sox4* and *Cd24a* expression over the AKI course, Mean ± SEM, one way ANOVA with Bonferroni, ** pValue<0.01 compared to Control. Scale: 4x – 2500*µ*m, 40x – 100*µ*m, 60x – 25*µ*m. Related to Figure S3, Table S2, S3.

Both scRNA-seq and RNAscope showed further kidney recovery at UIR Day 11 and 14, defined by restored *Slc34a1* expression, absent *Kim1*-positive injured proximal tubules and decreased immune infiltration (Figures S3D-S3F and 3A-3C). Injury resolution was confirmed with H&E staining (Figure S3A). The molecular analysis performed on all experimental time points showed the decrease of kidney injury markers Kim1 and Lcn2 over the course of AKI recovery, which returned back to normal by Day 14 (Figures S3B and S3C). While the small “Dedifferentiated Prox” cluster was still present at UIR Day 11, the kidney developmental signature was markedly weaker at this point (Table S2); these cells were not present at UIR Day 14. Similar to the Control kidney, UIR Day 14 proximal tubules were enriched with transmembrane transport and metabolic pathways, as the GO shows (Figure S3G), which reflects the normal proximal tubule homeostatic functions. These experiments examined multiple stages of AKI progression, thereby providing a powerful resource to dissect the gene expression changes in multiple cell populations involved in kidney injury and recovery.

Since proximal tubules are crucial contributors to the kidney injury response (Takaori et al., 2016), we further examined the gene expression changes characterizing the proximal tubular injury over the AKI course. We performed differential gene expression analysis of the “Injured Prox” compared to control at UIR Days 1, 2, 4 and 7, with a Venn diagram showing 183 common shared overlapping genes (Figure 3D). Noteworthy, the overlapping genes were enriched with pro-inflammatory pathways, such as leukocyte activation and degranulation (*Lcn2, Lgals1, Anxa2, S100a11, Cstb*) and viral process (*Kim1, Ifi27, Cd74, Anxa2*) (Table S3). Of particular interest, we identified the interferon-response gene *Ifi27*, encoding interferon alpha inducible protein 27, previously associated with drug-induced nephrotoxicity (Dieterich et al., 2009), among the genes shared by the “Injured Prox” of UIR Day 1, 2, 4 and 7 (Table S3). Moreover, we noticed the persistent interferon-response gene signature (*Ifit2, Ifit3, Ifitm3, Ifi44*) not only in the injured proximal tubules, but also in the collecting ducts and distal tubules (Table S1). Interferon signaling is known to play a role in renal inflammatory and autoimmune pathologies (Anders et al., 2010), and, as we and others (Wu et al., 2019) report, the renal epithelium acquires a pro-inflammatory phenotype in response to the enduring injury. Remarkably, we identified that renal developmental genes *Cd24a* and *Sox4* marked the injured proximal tubules throughout the AKI course, as the differential gene expression analysis of the UIR Day 1, 2, 4 and 7 “Injured Prox” UIR Day 1, 2, 4 and 7 shows (Table S3).

Next, we examined the gene expression changes occurring in the “Mixed Identity Cluster” over the AKI course. The differential gene expression analysis of the genes elevated in the mixed identity cells compared to control proximal tubules at UIR Days 1, 2 and 4 identified 99 genes common at all time points, as the Venn diagram shows (Figure 3E, Table S3). We identified *Umod* (loop of Henle) and *Aqp2* (collecting duct) among the overlapping genes, consistent with the mixed identity. GO analysis revealed elevation of cytokine response (*Ifi27, Ifitm3, Anxa2, Lcn2*), vesicle fusion (*Sparc, Tubb5, Tagln2, Lgals3*) and programmed cell death regulation (*Clu, Nfkbia, Ubb, Pea15a, Lgals1, Ctsd*) (Figure S3H). Importantly, we noticed the significant elevation of *Clusterin (Clu)*, in many injured kidney cell populations, including the “Mixed Identity Cells”, proximal tubules and collecting duct (Table S1). *Clu* expression might play a protective role in the kidney injury, since *Clu* deficiency results in worsened AKI outcome and elevated fibrosis (Guo et al., 2016). Previous work (Dieterle et al., 2010) reported urinary clusterin as a kidney injury marker, along with cystatin C (*Cst3*) and β2-microglobulin (*B2m*), two other AKI induced genes the scRNA-seq identified (Tables S1, S3). Importantly, we found that kidney developmental genes marked the “Mixed Identity Cells” over the AKI course (Figure S3H). Particularly, ToppCluster analysis of the genes shared by the “Mixed Identity Cells” at UIR Day 1, 2 and 4 (Kaimal et al., 2010) identified *Cd24a* and *Sox4* along with *Aqp2* (collecting duct) and *Umod* (loop of Henle) mentioned above (Figure S3I). pPCR performed on the independent animal cohort validated the injury inducible *Cd24a* and *Sox4* elevation shown by the scRNA-seq, which reversed by UIR Day 14, consistent with kidney recovery (Figure 3F). Overall, we demonstrated that renal developmental genes label renal epithelial injury, particularly injured proximal tubules and mixed identity cells, throughout the AKI course.

### *Sox4* and *Cd24a* Label the Proximal and Distal Tubule Injury

Then, we further examined the renal developmental gene expression in the injured kidney. As scRNA-seq found and qPCR validated, *Sox4* was elevated in the adult kidney at multiple AKI stages. Previous work shows that *Sox4* plays a crucial role in multiple developmental processes, including kidney, heart and immune system development (Hu and Chen, 2013; Huang et al., 2013; Paul et al., 2014; She and Yang, 2015). *Sox4* was implicated in numerous human malignancies (Bilir et al., 2016; Tavazoie et al., 2008); moreover, the recent study reported that *Sox4* promotes renal cell carcinoma metastasis via TGFβ-induced EMT (Ruan et al., 2017). However, the role of *Sox4* in AKI is unknown.

The scRNAseq data indicates the broad AKI induced *Sox4* upregulation in multiple nephron tubule segments (Figure 2F, Table S1). Examining the pattern of *Sox4* expression revealed a strong inverse relationship with *Slc34a1*-positive proximal tubules at UIR Day 4, when the injury starts resolving, as the combined feature plot (Figure 4A). Thus, we questioned whether *Sox4* specifically marks the injured proximal tubules during the AKI response. To test this, we hybridized specific *Sox4, Slc34a1* and *Kim1* RNAscope probes to the injured and control kidney sections and found enhanced *Sox4* expression in the *Kim1*-positive injured proximal tubules, exhibiting weakened *Slc34a1* signal, at UIR Day 1 and 2 (Figure S4A). As previously described, UIR Day 4 and 7 exhibit substantially reduced *Kim1* and restored *Slc34a1* expression, consistent with kidney recovery. Notably, *Sox4* was predominantly confined to remaining injured *Kim1*-positive proximal tubules at UIR Day 4 and 7 (Figures 4B, S4A), thus validating the scRNA-seq prediction. Quantitative IMARIS analysis revealed a substantial difference between *Sox4* transcript number in the *Kim1*-enriched injured vs *Slc34a1*-enriched mature proximal tubules (Figure S4B). Thus, we showed that *Sox4* is a marker of proximal tubule dedifferentiation in adult AKI.

**Figure 4.**
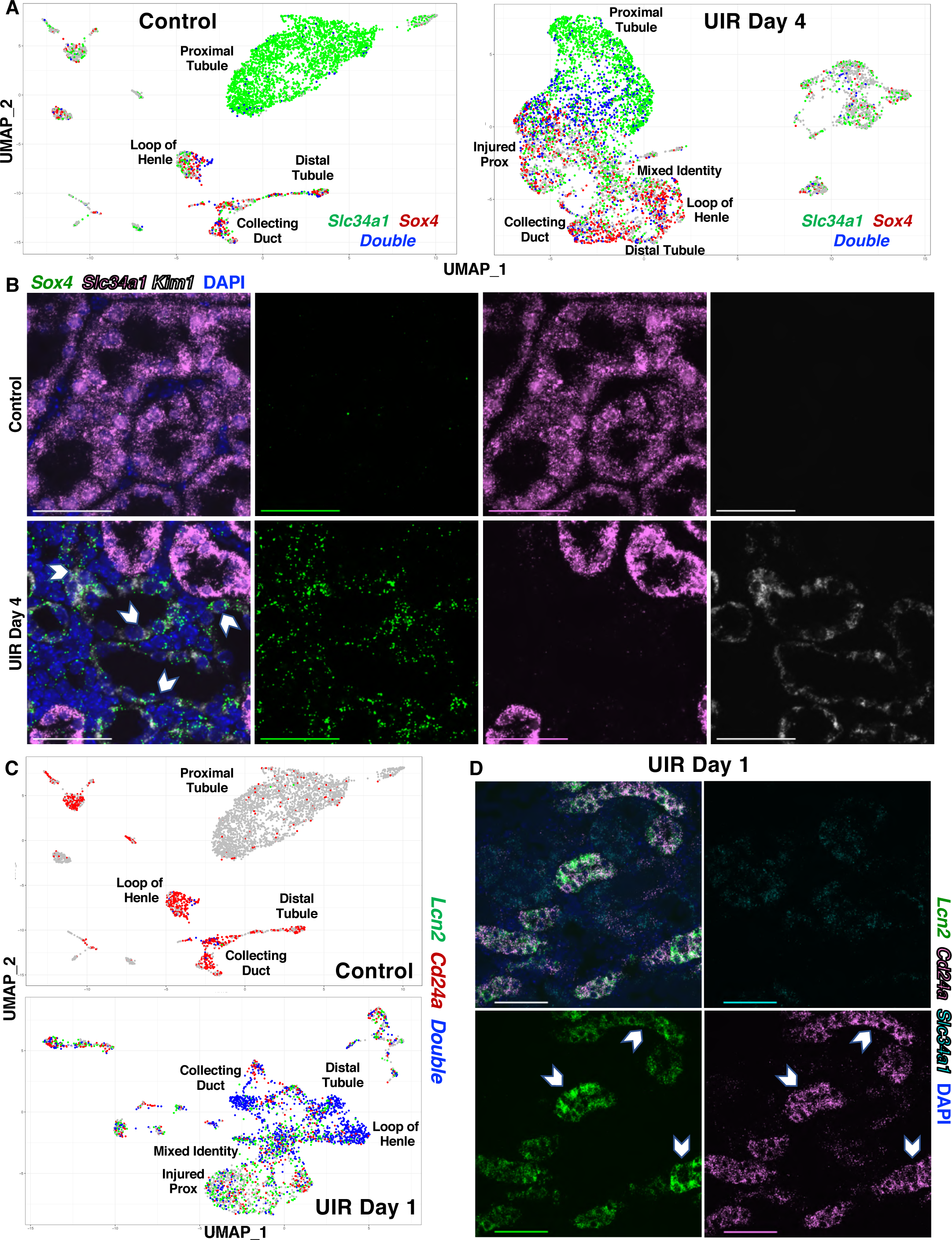
*Sox4* and *Cd24a* Label the Proximal and Distal Tubule Injury. (A) Feature plots show *Slc34a1* (green) and *Sox4* (red) and double (blue), expression in UIR Day 4 vs Control. (B) RNAscope with *Slc34a1* (pink), *Sox4* (green) and *Kim1* (white) probes, DAPI (blue), UIR Day 4 vs Control. 60x Nyquist zoom, 0.14 *µ*m/px, Maximal Intensity Projection (MaxIP) from Z-stack, scale 50 *µ*m. (C) Feature plots show *Lcn2* (green), *Cd24a* (red) and *Double* (blue) positive cells in UIR Day 1 vs Control. (D) 60x RNAscope images show *Lcn2* (green) and *Cd24a* (pink) colocalization at UIR Day 1, 60x Nyquist zoom, 0.21 *µ*m/px, Maximal Intensity Projection (MaxIP) from Z-stack, scale 50*µ*m. Related to Figures S4 and S5.

Next, we assessed whether renal developmental genes mark renal injury in the distal nephron tubule segments. We observed that *Cd24a* notably overlaps with *Lipocalin 2* (*Lcn2)*, a clinically recognized distal marker (Devarajan, 2008, 2010; Mishra et al., 2005) used in clinical practice as an early predictor and progression marker of AKI (Khawaja et al., 2019; Sun et al., 2018; Zhang et al., 2017a). As the combination UMAPs show, the injured kidney exhibits a prominent number of double *Cd24a; Lcn2* positive cells in the loop of Henle, distal tubule, collecting duct and “Mixed Identity Cells” at UIR Day 1 (Figure 4C). We detected the highest *Lcn2* and *Cd24a* levels at the most severe stages of kidney injury (UIR Day 1 and 2), which returned back to normal by Day 14 (Figures 3F, S3B, S3C). Examining the spatial pattern of *Cd24a* and *Lcn2* expression with RNAscope revealed their striking level of colocalization at multiple time points after AKI (Figure 4D, S5A).

The scRNAseq data also showed that *Cd24a* increases in the proximal tubules in response to the injury, as discussed earlier. To quantitatively assess the *Cd24a* expression in the different injured kidney compartments, we counted *Cd24a* within *Slc34a1*-positive proximal tubules vs *Lcn2*-enriched injured tubules using IMARIS. We found that UIR caused *Cd24a* increase in *Slc34a1*-positive proximal tubules at Day 1, 2 and 4, however, it was substantially more enriched in the *Lcn2*-positive injured compartment (Figures S5B, S5C). Moreover, we found that *Cd24a* and *Lcn2* transcript numbers exhibit remarkable positive correlation, as the Pearson analysis shows (Figure S5D). Importantly, AKI induced *Cd24a* transcriptional changes were reproduced on the protein level, as we show with Cd24a immunohistochemistry (IHC) (Figure S5E). Thus, we demonstrate that renal developmental genes might serve as new AKI markers.

### Novel Gene Expression Signatures of AKI

With scRNA-seq, we found that AKI induced some global gene expression changes in the kidney, in multiple compartments, exemplified by the cytokine and *Clusterin* signaling discussed earlier. We also found the widespread *Spp1* (a.k.a. *Osteopontin*) upregulation caused by AKI, including in tubular, stromal and immune cells (Figures 5A and 5B). Spp1 is a secreted phosphoprotein 1 which plays a major role in bone metabolism and immune system activation, particularly, macrophages and T- and B-cell mediated immunity; Spp1 is implicated in multiple renal pathologies, including renal cell carcinoma, diabetic nephropathy and allograft rejection (Kaleta, 2019). Consistent with the previous work, we observed that *Spp1* exhibits some baseline expression mostly in the renal medulla; however, UIR caused striking *Spp1* elevation in both cortex and medulla (Figure 5A). We observed that *Spp1* expression overlaps with another kidney injury associated gene, *Krt8*, in the injured nephron tubules and collecting duct (Figure 5B). Elevated cytokeratin tubular expression is a known characteristic of epithelial cell stress (Djudjaj et al., 2016; Liu et al., 2017); consistent with these observations, we identified Krt7, 8 and 18 elevation in the UIR injured tubules (Table S1). Notably, aberrant cytokeratin expression is associated with tumor progression and metastasis (Fang et al., 2017). In particular, Krt8 was shown to play a crucial role in renal cell carcinoma (Tan et al., 2017). Immunofluorescence (IF) showed that *Krt8* transcript elevation was accompanied by increased protein (Figure S6A); moreover, we identified Krt8-positive renal tubules which also expressed *Spp1*, using a combined CISH and IF approach (Figure 5C). We observed that Krt8 and *Spp1* levels remained elevated through UIR Day 4 and lowered to the normal levels by Day 14, indicating injury resolution (Figure S6A). Overall the results show elevation of multiple established kidney epithelial injury markers and reveal the novel AKI induced overlapping *Spp1* and *Krt8* expression. Since both *Spp1* and *Krt8* play a role in renal cell carcinoma metastasis and progression, dual *Spp1-* and *Krt8-* positive cells give evidence for renal tubular cells acquiring a more mesenchymal phenotype in response to the injury.

**Figure 5.**
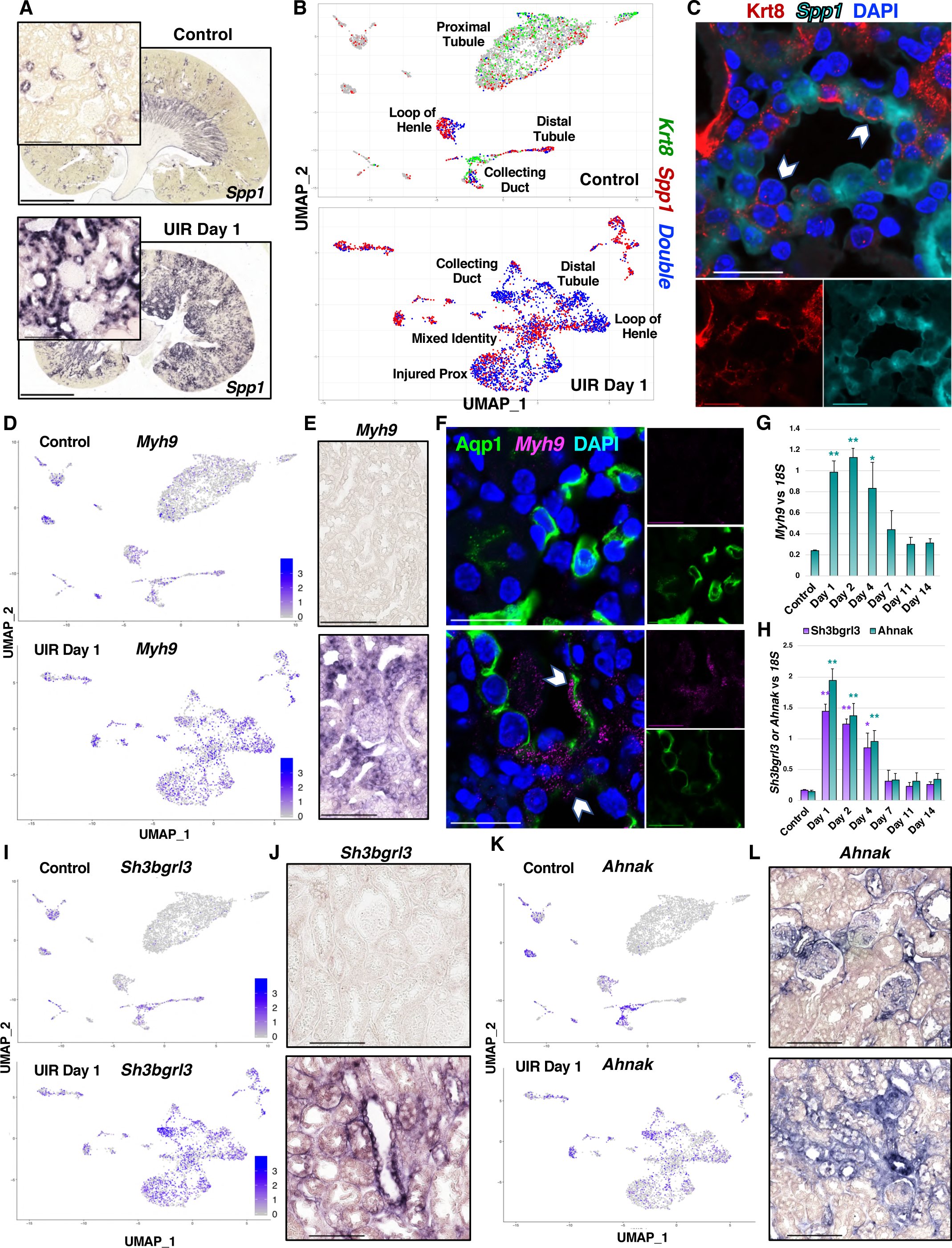
Novel Gene Expression Signatures of AKI. (A) *Spp1* CISH, Control vs UIR Day 1, 4x and 40x zoom into the cortex. (B) Combined Feature plots show *Krt8* (green), *Spp1* (red) and *Double* (blue) positive cells in UIR Day 1 vs Control. (C) *Spp1* CISH (cyan), Krt8 IF (red), DAPI (blue), UIR Day 1, 60x Nyquist 3x zoom. Pointers show *Spp1* and Krt8 overlap. The combined scRNA-seq and CISH results define the precise cell types with elevated *Spp1* and *Krt8* elevated expression following UIR. (D) Feature plots show *Myh9* expression in UIR Day 1 vs Control. (E) *Myh9* CISH shows elevated expression in UIR Day 1 tubules, glomeruli and interstitium compared to the Control, 40x. (F) *Myh9* CISH (magenta), Aqp1 IF (green), DAPI (blue), Control vs UIR Day 1, 60x Nyquist 3x zoom. Pointers show *Myh9* within Aqp1-positive tubules. (G and H) qPCR shows *Myh9, Sh3bgrl3* and *Ahnak* expression over AKI course, Mean + SEM, 4-6 animals per group, one way ANOVA with Bonferroni and Holm, * p<0.05, ** p<0.01 compared to the Control. (I) Feature plots how *Sh3bgrl3* expression in Control vs UIR Day 1. (J) *Sh3bgrl3* CISH shows elevated expression in UIR Day 1 tubules and interstitium compared to the Control, 40x. (K) Feature plots show *Ahnak* expression in Control vs UIR Day 1. (L) *Ahnak* CISH shows expression in the glomeruli and interstitium of the Control, and elevated expression in UIR Day 1 tubules and interstitium, 40x. Scale 4x, 2500um; 40x, 100um; 60x Nyquist 3x zoom, 25*µ*m. Related to Figure S6.

scRNAseq also identified the elevation of *Myosin Heavy Chain 9*, or *Myh9*, a non-muscle myosin involved in cell motility, cell shape maintenance and cytokinesis, in the injured renal tubules. As we and the others (Arrondel et al., 2002) show, normal adult kidney mostly expresses *Myh9* in the stromal cells, pericytes, glomeruli and endothelial cells, with very faint tubular expression (Figure 5D). As both scRNAseq and CISH indicated, UIR caused a marked *Myh9* elevation in all renal tubule compartments, including the sub-clusters of injured and dedifferentiated proximal tubules, and in the collecting ducts (Figures 5D and 5E). Importantly, previous work specifically reported that Myh9 expression is absent in normal proximal tubules (Otterpohl et al., 2017). We validated the injured proximal tubule expression via co-staining the *Myh9* CISH with Aqp1 IF and identified Aqp1-positive proximal tubules exhibiting elevated *Myh9* expression which was not detected in the control (Figure 5F), providing an additional evidence for the injury induced mesenchymal phenotype in renal epithelial tubules. With qPCR, we show that *Myh9* elevation persists to UIR Day 4 and then reverses to the normal levels (Figure 5G).

The scRNAseq approach allowed us to identify additional novel genes not previously implicated in AKI. For examples, we found that *Sh3bgrl3* and *Ahnak* were upregulated in the UIR treated kidneys, as validated by qPCR (Figure 5H). *Sh3bgrl3* encodes the SH3 Domain-Binding Protein 1, associated with GTPase, oxidoreductase and anti-apoptotic activity (Henn et al., 2001). The control kidney exhibited very low levels of *Sh3bgrl3*, while it was highly enriched in all tubular lineages in UIR Day 1 (Figure 5I). CISH confirmed the AKI induced *Sh3bgrl3* elevation within renal tubules and in peritubular spaces (Figure 5J). The other gene, *Ahnak*, encodes Neuroblast Differentiation-Associated Protein AHNAK, involved in pathogenesis of Miyoshi Muscular Dystrophy (Cacciottolo et al., 2011) and tumor metastasis via TGF-β induced epithelial-to-mesenchymal transition (Sohn et al., 2018). Notably, both scRNAseq and CISH showed that *Ahnak* is present in the Control podocytes and stroma with some tubular expression (Figures 5K and 5L), however, UIR resulted in the marked *Ahnak* elevation in the tubular and stromal compartments. With qPCR, we showed that both *Ahnak* and *Sh3bgrl3* stay elevated until UIR Day 4, and then return to normal expression levels (Figures 5G, 5H). As the scRNAseq data shows, both *Ahnak* and *Sh3bgrl3* are highly enriched in the injured proximal tubules while nearly absent in the healthy ones, suggesting they represent novel proximal tubule injury markers.

### scRNAseq Identifies Novel Epithelial-to-Stromal Interactions in the Adult AKI

Growing evidence suggests the importance of cell-to-cell interactions in kidney injury progression (Humphreys, 2018). Thus, we examined the molecular nature of normal and pathologic intracellular interactions in the kidney. To this end we paired the cells enriched with ligand to the cells enriched for the corresponding receptor, and found, for examples, that the proximal and distal tubules and the stromal cells might interact with each other via the calmodulin (*Calm1, Calm2, Calm3*), growth factor (*Egf, Fgf1, Hbegf, Vegfa, Mdk, Igfbp4*) and lipid metabolism (*Lpl, Lrpap1, Psap*) pathways in the normal kidney (Figure 6A, Table S4). We also found that the stromal cells might influence proximal tubules via collagen signaling, since the stromal cells were enriched with *Col1a1/1a2, Col3a1, Col4a1/4a5* and *Col18a1*, while the proximal tubules expressed their receptors. Our data also highlights potential pericyte-to-proximal tubule interactions via the collagen, nidogen (*Nid1, Nid2*), matrix metalloproteinase (*Timp2*) and calmodulin signaling.

**Figure 6.**
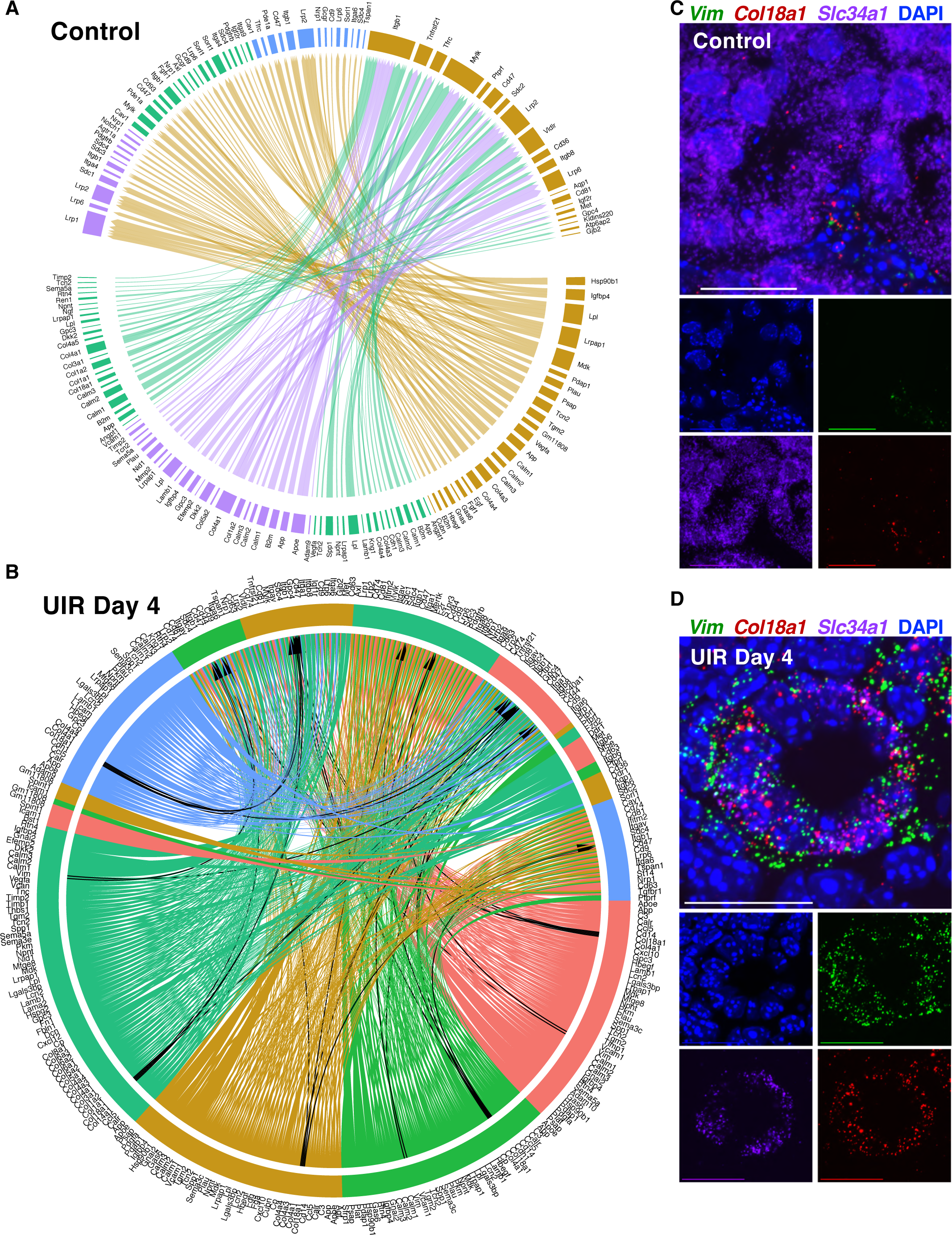
scRNAseq Identifies Novel Epithelial-to-Stromal Interactions in the Adult AKI. (A) Circos Plot of ligand-receptor interactions between the proximal (brown) and distal (blue) tubules, the stromal cells (green), and the stromal/pericyte cells (purple) in the normal kidney. (B) Circos Plot of ligand-receptor interactions between the proximal tubules (brown), injured prox (red), mixed identity cells (green), distal tubules (blue) and the stromal cells (teal) in the normal kidney. Black arrows show *Vim-Cd44*, *Col18a1-Gpc4* and *Col18a1-Itgb1* Ligand-Receptor pairs. Note the dramatic increase in the number of potential interactions compared to control, using the same filters. (C) Representative RNAscope image of Control proximal tubules, *Vim* (green), *Col18a1* (red), *Slc34a1* (purple), 60x 4x Nyquist zoom, MaxIP from the Z-stack. (D) Representative RNAscope image of UIR Day 4 proximal tubule, *Vim* (green), *Col18a1* (red), *Slc34a1* (purple). 60x 4x Nyquist zoom, MaxIP from the Z-stack. Scale 50*µ*m. Related to Figure S6, Table S4, Movie S6 and S7.

Next, we examined how AKI affects cell-to-cell communication in the kidney and focused on UIR Day 4, as an intermediate injury response time point. Ligand-to-receptor analysis revealed that AKI causes a dramatic increase in the interactions between the renal cell populations. In particular there was elevated extracellular matrix signaling by the injured epithelial cells (Figure 6B, Table S4). We observed the elevation of *Vimentin (Vim)*, which is known as an activated fibroblasts marker associated with EMT (Liu et al., 2015; Liu, 2010) and kidney fibrosis (Grande et al., 2015; Humphreys, 2018), in the injured proximal tubules and mixed identity cells (Figure S6B). Moreover, we found that the Vim receptor encoding gene *Cd44* (Pall et al., 2011) was expressed in the stromal cells, suggesting the pathologic injury induced tubular-to-stromal crosstalk. Importantly, we noticed that *Cd44* was also enriched in the injured proximal tubules and in the mixed identity population themselves, suggesting that injured renal epithelial cells not only interact with stromal cells, but crosstalk with each other in the injury setting.

Of particular interest, we also observed that AKI induced elevation of *Col18a1*, encoding an extracellular nonfibrillar collagen which is a necessary component of basement membrane (Kinnunen et al., 2011). We and others (Harding et al., 2011; McMahon et al., 2008) show that *Col18a1* is normally present mostly in the glomerulus with some weak expression in the collecting duct (Figure S6B), however, its role in the AKI response remains unknown. We found remarkable *Col18a1* elevation in the injured proximal and distal tubules and the mixed identity cells caused by AKI; moreover, our analysis identified *Col18a1* receptors encoding genes *Gpc4* and *Itgb1* to be enriched in the stromal cells, which highlights another potential epithelial-to-stromal cell interaction pathway (Figure 6B, Table S4). These findings were validated by the qPCR, which showed significant *Vim* and *Col18a1* elevation caused by AKI (Figures S6C-S6D). Moreover, RNAscope revealed the marked *Vim* and *Col18a1* upregulation within the *Slc34a1*-positive proximal tubules at UIR Day 1, 2 and 4, which returned to the normal interstitial and periglomerular expression pattern by Day 14 (Figures 6C and 6D, S6E, Movies S5 and S6). Other factors involved in injury induced epithelial-to-stromal crosstalk included collagen (*Col4a1, Col4a3, Col4a4*), lipocalin 2 (*Lcn2*), interferon (*Ifitm2*), galectin (*Lgals3bp*) and Osteopontin (*Spp1*) (Table S4). Thus, our scRNAseq experiments revealed that renal epithelial and stromal cells show increased potential interactions with each other in the condition of injury.

### Renal Developmental Genes Outline Kidney Fibrotic Remodeling

Next, we tested if the AKI induced genes identified by the scRNAseq could serve as markers of maladaptive injury response and fibrotic remodeling. We induced the identical UIR in 10 weeks old male Swiss-Webster mice (Figure 7A) and found that they exhibited significantly elevated fibrosis marker expression, prominent fibrotic remodeling and decreased mature proximal tubule marker *Slc34a1* expression at UIR Day 14 compared to the younger mice (Figures 7B–7D). Thus, we show that age of AKI onset is a definitive factor in determining whether the kidney undergoes repair or maladaptive remodeling. Notably, we observed that both *Sox4* and *Cd24a* were significantly elevated for longer times in the older animals compared to the younger ones (Figures 7E, 7F). Moreover, IHC showed that *Cd24a* stays elevated in renal tubules of the older mice throughout the AKI course, while it resolves in the younger animals by UIR Day 14 (Figure 7G). This shows that AKI induced *Cd24a* transcriptional increase also occurs at the protein level. Overall, our data reveals that renal developmental genes outline AKI progression and might serve as predictors of the injury outcome.

**Figure 7.**
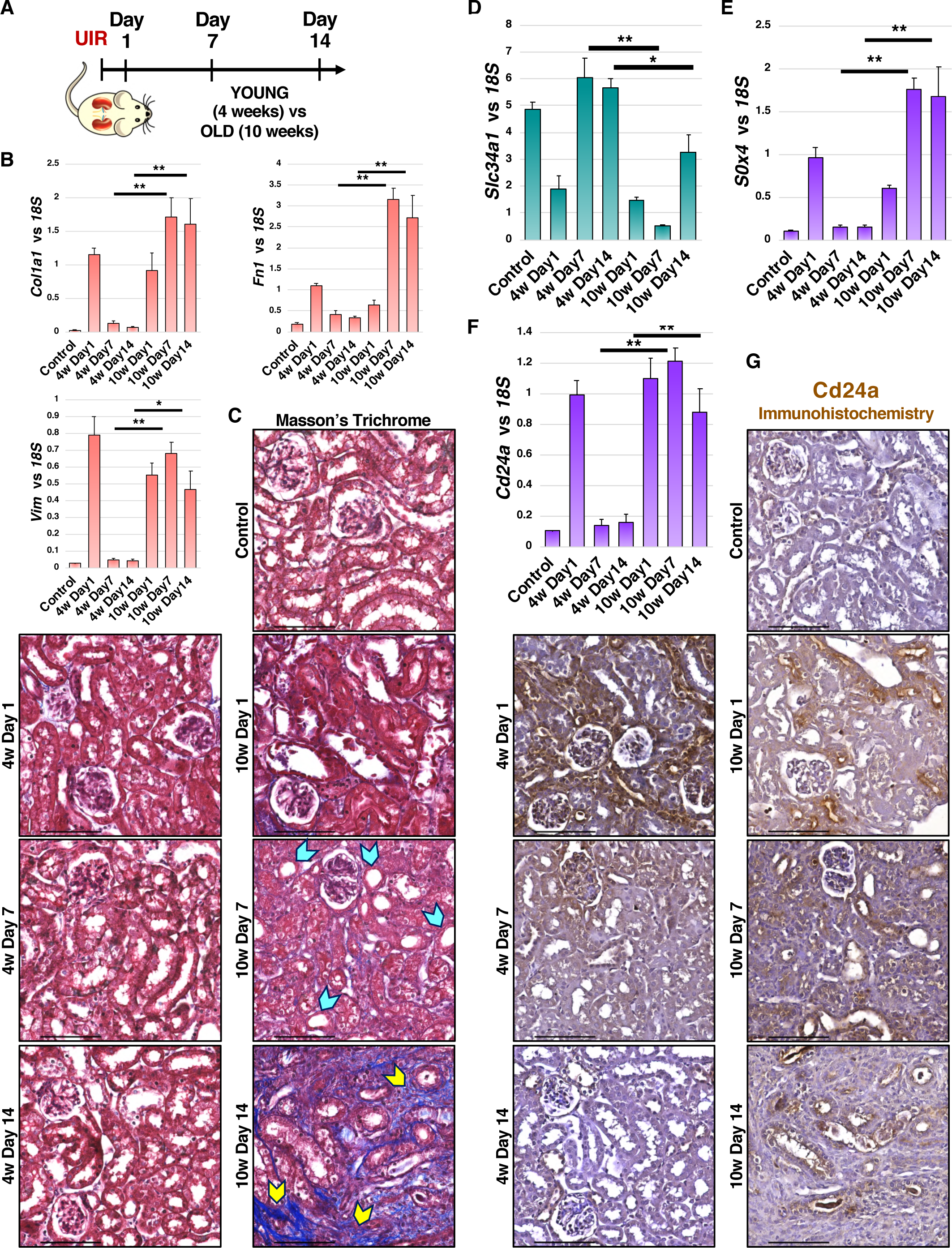
Renal Developmental Genes are Elevated in Kidney Fibrotic Remodeling. (A) Experimental outline. (B) qPCR shows the fibrosis markers *Vim, Col1a1* and *Fn1* expression over the AKI course, Mean ± SEM, n=4-6 per group, Student’s *t* test, 4w Day 7 vs 10w Day7, 4w Day 14 vs 10w Day 14, * p<0.05, ** p<0.01. (C) Masson’s Trichrome staining shows fibrotic remodeling in the 10w old mice at UIR Day 14; tubular dilation - blue pointers, fibrotic areas - yellow pointers; 40x. (D-F) qPCR shows *Slc34a1, Sox4* and *Cd24a* expression over the AKI course, Mean ± SEM, n=4-6 per group, Student’s *t* test, Day 7 4w vs 10w, Day 14 4w vs 10w, * p<0.05, ** p<0.01. (G) Cd24a IHC, 40x. Scale 100*µ*m.

## Acknowledgements

The authors thank members of Jiang and Yutzey laboratories for chromogenic *in situ* hybridization and RNAscope protocols. The authors thank members of Cincinnati Children’s Hospital Medical Center Gene Expression Core, Confocal Imaging Core and Veterinary Services Surgical Core for valuable guidance and assistance with imaging, scRNA sequencing and animal procedures. This work was supported by NIH grant DK120842 (SSP), Mallinckrodt Fellowship Award (KAD).

## Author Contributions

V.R-M., M.A. and S.S.P. prepared the manuscript. V.R-M. assembled figures. A.P. did kidney dissociation and scRNA-seq. M.A. analyzed the scRNA-seq data. Q.M. and V.R-M. induced UIR. V.R., M.A., P.D. and S.S.P. developed experimental strategy and analyzed results. V.R-M., S.M.C. and Q.M. did validation experiments. M.P.S. and J.M.K. designed imaging configurations and provided guidance for RNAscope data analysis. K.A.D. helped to cover the sequencing costs.

## Methods

### Lead Contact and Materials Availability

Further information and requests for resources and reagents should be directed to and will be fulfilled by the Lead Contact, Dr. S. Steven Potter.

### Experimental Model and Subject Details

The Institutional Care and Use Committee (IACUC) of Cincinnati Children’s Hospital Medical Center reviewed and approved all animal procedures used in this study. The unilateral ischemia/reperfusion (UIR) (Le Clef et al., 2016; Yang et al., 2010) was induced via atraumatic left renal pedicle clamping for 30 min at 37°C in the 4 weeks old male Swiss-Webster mice, and the kidneys were harvested at Day 1, 2, 4, 7, 11 and 14 after the UIR (n=1 per each time point) for the single cell RNA sequencing (scRNAseq) and histological verification. Kidneys harvested from the naïve mice of the same strain and age were used as controls. The validation was performed on the additional set of animals treated with the identical UIR procedure (n=3-6 per group). For examining the effects of age on the AKI outcome, the equivalent UIR was induced in the 10 weeks old male Swiss-Webster mice, and the kidneys were harvested at UIR Day 1, 7 and 14 (n=4 per group). All animal procedures were in concordance with the institutional protocols.

### Method Details

#### Experimental Animals and AKI Procedure

All animal procedures were conducted according to the Institutional Care and Use Committees (IACUC) of Cincinnati Children’s Hospital Medical Center guidelines. Swiss-Webster (CFW) 4 and 10 weeks old male mice obtained from Charles River Laboratories were subjected to the UIR model of acute kidney injury (AKI). The animals were anesthetized with isoflurane and placed on the homeothermic blanket, and the body temperature was maintained at 37°C. Ischemia was induced via the midline abdominal approach with the atraumatic vascular clamp placed on the left kidney pedicle and verified with the immediate kidney color change (Skrypnyk et al., 2013), with the right kidney left undisturbed. The clamp was released after 30 min followed by the left kidney color change back to red. The animals were closed via abdominal muscle and skin suturing, given 1ml of warm sterile normal saline subcutaneously for the fluid loss compensation and returned to the normal housing. The animals received 50ml of 0.03mg/ml Buprenorphine subcutaneously for analgesia on the day of surgery and on the next day, and were monitored daily according to the IACUC requirements.

#### Single Cell Suspension Preparation and scRNA-seq procedure

The 4 weeks old male CFW mice treated with the UIR procedure were intraperitoneally injected with 100 μL heparin (100 U/mL), anesthetized with isoflurane chamber and euthanized via exsanguination followed by cervical dislocation at Day 1, 2, 4, 7, 11 and 14 after the UIR (n=1 per group). The kidneys were perfused with ice-cold DPBS via the aorta prior to the harvesting to remove the red blood cells, and the left (injured) and the right (contralateral) kidneys from the UIR treated mice were collected along with the control kidneys obtained from the naïve mice. The kidneys were placed on ice-cold Dulbecco’s phosphate-buffered saline (DPBS), decapsulated, bisected lengthwise and sliced coronally. The cortical regions were then finely minced with a sterile razor blade until tissue was homogenous and clumps were broken down, and 65 mg of the minced tissue was placed in 2 ml digestion buffer containing 3 mg/mL Type 2 collagenase (Worthington, Collagenase Type 2), 1.5 mg/mL ProNase E (Sigma P6911), 62.5 U/mL DNAse (Applichem, A3778), and 5 mM CaCl_2_ made up in DPBS. The digest mix was incubated in a 37 °C water bath for 20 minutes with vigorous trituration with a 1 mL pipet every 2 min. 5 *µ*L aliquots were taken every 5 min and visualized using scope to ensure adequate digestion. After 20 min, the digest mix was added to two 1.5 mL low-adhesion tubes and was then incubated in a thermomixer for 5 min at 37 °C while shaking at 1400 RPM. The digest mix was subsequently added to a 70 *µ*M filter (Miltenyi, 130-098-462) stacked on a 30 *µ*M filter (Miltenyi, 130-098-458) on a 50 mL conical tube. The filters were rinsed with 10 mL PBS/BSA 0.01% and the flow-through was then transferred to a 15 mL conical tube. The cells were pelleted by centrifugation at 350 g for 5 min at 4 °C. The supernatant was discarded, and the pellet re-suspended in 100 *µ*L PBS/BSA. The cells were visualized and if >30% were RBC, RBC lysis step was performed: 1 mL of RBC lysis buffer (Sigma, R7757) was added, and the cells were triturated 20x with a 1 mL pipet; the cells were then incubated for 2 min on ice. 12 mL ice-cold PBS/BSA was then added to dilute the RBC lysis buffer. If no RBC lysis was necessary, then 12 mL ice-cold PBS/BSA was added to the cells. The cells were pelleted by centrifugation at 350 g for 5 min at 4 °C. The supernatant was then discarded, and the cells re-suspended in 2 mL PBS/BSA. Cells were analyzed with a hemocytometer using trypan blue and the concentration was adjusted to 100,000 cells/mL for the droplet-based single-cell RNA-seq procedure (DropSeq) based on a protocol from Macosko et. al (Macosko et al., 2015).

The remaining tissue slices from each kidney were fixed with 4% paraformaldehyde (PFA) in PBS overnight (O/N) at 4°C and paraffin embedded for histological assessment and snap-frozen in liquid nitrogen for the molecular analysis. The validation animals (n=3-4 per group) treated with the identical UIR protocol were harvested at the same time points (Day 1, 2, 4, 7, 11 and 14), and the kidney tissue was fixed in 4% PFA O/N at 4°C and snap-frozen in liquid nitrogen for the further analysis.

#### scRNAseq Data Analysis

The generated cDNA libraries were quantified using an hsDNA chip and were sequenced on the Illumina HiSeq 2500 using one flow cell (about 300 million reads) per sample. The raw fastqs were processed by aligning read2 to the mm10 genome using bowtie2-2.2.7 using the -k 1 option. The aligned reads were tagged with their corresponding UMI and barcode from read1. Each aligned read was tagged with its gene name. An expression matrix was generated by counting the number of unique UMIs per gene per cell.

Cell-type clusters and markers genes were identified using the R v3.6.1 library Seurat v3.1.0 (Butler et al., 2018; Stuart et al., 2019). All clustering was unsupervised, without driver genes. The influence of the number of unique molecular identifiers was minimized by regression within the ScaleData function. Initial cell filtering selected cells that expressed >500 genes. Genes included in the analysis were expressed in a minimum of three cells. Only one read per cell was needed for a gene to be counted as expressed per cell. The resulting gene expression matrix was normalized to 10,000 molecules per cell and log transformed (Macosko et al., 2015). Cells containing high percentages of mitochondrial, >30%, and hemoglobin genes, >0.025% were filtered out. Genes with the highest variability among cells were used for principal components analysis. Cell clusters were determined by the Louvain algorithm. Dimension reduction was performed using the Python implementation of UMAP (Uniform Manifold Approximation and Projection) using significant principal components determined by JackStraw plot. Marker genes were determined for each cluster using the Wilcoxon Rank Sum test within the FindAllMarkers function using genes expressed in a minimum of 10% of cells and fold change threshold of 1.3. Over/under clustering was verified via gene expression heatmaps.

Putative signaling interactions between proximal tubules, injured proximal tubules, stromal cells, and mixed-identity cells were assessed. Potential receptor-ligand interactions were found by pairing a cell-type expressing a ligand with a cell-type expressing its receptor pair. A receptor or ligand was considered expressed in a cell-type having an average normalized expression of >0.25. Receptor-ligand pairs were determined using the curated receptor-ligand database by the RIKEN FANTOM5 project (Lizio et al., 2015). Receptor-ligand pairings for each cell type were visualized by a chord diagram using the R package circlize (Gu et al., 2014).

Differential gene expression between control proximal tubules, injured proximal tubules and “Mixed Identity Cells” was determined by Wilcoxon Rank Sum test within Seurat’s FindMarkers function using a log fold change threshold of 0.5 and an adjusted p-value cutoff of 0.01. Venn diagrams were prepared with the R package VennDiagram (Chen and Boutros, 2011).

### RT-qPCR

Gene expression changes identified by scRNA-seq were validated using the Real-Time Quantitative PCR. Three to six mice from the original scRNA-seq cohort and the additional validation cohort were used per each group. Total RNA was isolated from the homogenized whole kidney lysates with RNA Stat-60 extraction reagent (Amsbio, CS-111) and purified using the GeneJET RNA purification kit (Thermo Fisher Scientific, KO732). The cDNA was synthesized with the iScript Reverse Transcription Supermix (Bio-Rad, 1708841). qPCR was performed with TaqMan universal PCR master mix (Thermo Fisher Scientific, 4304437) on the Applied Biosystems Quant Studio 3 RT PCR system. The TaqMan primers details are presented in the Table S5. The reported Ct values are the mean of two replicates of the same cDNA sample. The target gene Ct values were normalized to the eukaryotic 18S rRNA endogenous control and presented as the fold change.

### Western Blotting

We validated the RNA expression changes detected by the scRNAseq with Western blots. Three to six mice from the original scRNA-seq cohort and the additional validation cohort were used per each group. Total protein was extracted from the homogenized whole kidney lysates using the M-PER mammalian protein extraction reagent (Thermo Fisher Scientific, 78501) supplemented with protease (Thermo Fisher Scientific, 78430) and phosphatase (10 mM Sodium Orthovanadate) inhibitors. 10 *µ*g of protein was separated on 4-12% polyacrylamide gel by SDS electrophoresis, transferred to PVDF membrane, blocked with the Odyssey blocking buffer (LI-COR, 927-40000) and incubated with the target recognizing primary antibodies (Kim1, goat, R&D Systems, AF1817, 0.25 *µ*g/ml; Lcn2, rabbit, MBL, JM-3819-100, 0.5 *µ*g/ml) O/N at 4 °C. On the next day, the membranes were washed with Tris buffered saline 0.02% Tween (TBST) and incubated with the secondary fluorescent antibodies for 1 hour at RT in the darkness. The target protein levels were normalized to the endogenous control (GAPDH, mouse, Millipore Sigma, MAB374, 1/500). The membranes were scanned on the LI-COR Odyssey CLx imaging system.

### Chromogenic *In Situ* Hybridization (CISH)

CISH riboprobes were generated from the gDNA or cDNA with PCR and labeled with digoxigenin (DIG) using DIG RNA labeling mix (Roche, 11277073910) by *in vitro* transcription, and the primer sequences are listed in the Table S5. Primer specificity was checked with BLAST analysis. PFA fixed paraffin embedded (PFPE) 6 *µ*m kidney sections were subjected to the CISH protocol as previously described (Lan et al., 2001; Xu et al., 2016). On day 1, deparaffinated and dehydrated sections were incubated with Proteinase K (1ug/ml) for 30 min at 37 °C, fixed in 4% PFA for 30 min at room temperature (RT), acetylated in 0.25% acetic anhydride and hybridized for 14-16 hours at 70 °C with the antisense riboprobes. Sense riboprobes were used for the negative controls. All day 1 procedures were performed in the RNase-free conditions. On the second day, sections underwent a series of saline sodium citrate buffer washes and were incubated with the anti-DIG-AP fab fragments (Roche, 11093274910, 1:1000) O/N at 4 °C, followed by a series of post-hybridization washes and color development with BM Purple (Roche, 11442074001). Color development time varied from several hours to 3-4 days depending on the target gene expression levels. For the immunofluorescent (IF) co-labeling, the protocol was modified by adding the primary antibody recognizing the target (Krt8, rabbit, Abcam, EP1628Y, 1/100; Aqp1, mouse, Santa Cruz, sc-25287, 1/50) to the anti-DIG-AP antibody mixture and incubation O/N at 4 °C (Lopez, 2014). Next day, the sections were incubated with the fluorescent secondary antibodies for 1 hour at RT in darkness, followed by the post-hybridization washes and color development, according to the CISH protocol. DAPI (Thermo Fisher Scientific, 62248, 1:1000) treatment for nuclei labeling was performed after the color development; the sections were mounted with Vectashield antifade mounting medium (Vector Laboratories, H-1000). For the double CISH/IF staining, treatment with the antisense riboprobe and secondary antibodies only was used as the negative control. For the immunohistochemistry (IHC) staining, the sections were subjected to the endogenous peroxidase quenching, blocking and primary antibody incubation (Cd24a, rat, Abcam, ab64064, 1/100) O/N at 4 °C, followed by the signal detection with the ImmPRESS reagent kit with DAB substrate (Chihanga et al., 2018). CISH and IHC images were obtained on the Nikon Ti2 wide-field microscope. RGB images of CISH/IHC were taken with an Andor Zyla 4.2 plus camera and a Lumencor LIDA RGB transmitted light source.

### Single Molecule Fluorescent *In Situ* Hybridization (smFISH, using RNAscope)

RNAscope probes were purchased from Advanced Cell Diagnostics, Icn (ACD) and are summarized in the Table S5. RNAscope was performed with the Multiplex Fluorescent v2 assay (ACD, 323100) on the freshly sectioned PFPE 6 *µ*m kidney sections according to the manufacturer’s protocol. Briefly, the deparaffinized and dehydrated kidney sections underwent the endogenous peroxidase quenching, heat-induced target retrieval and protease digestion, followed by the incubation with up to three target riboprobes for 2 hours at 40 °C. All the aforementioned steps were performed in the RNase-free conditions. Next, tyramide signal amplification was performed according to the manufacturer’s protocol and conjugated to an Opal dye (PerkinElmer). The sections were treated with DAPI (Thermo Fisher Scientific, 62248, 1:1000) and mounted with Vectashield antifade mounting medium (Vector Laboratories, H-1000). The negative controls were treated with the signal amplification reagents but no target riboprobes and processed alongside the experimental sections.

### Fluorescent Microscopy and Quantitative Image Analysis

The double CISH/IF imaging was performed on a Nikon A1R HD confocal TiE microscope using the galvanometric scanner. The CISH signal was captured both using a transmitted light detector to detect absorption of photons by the chromogenic substrate and as near-IR fluorescence of the BM purple using a 647nm laser for excitation and capturing fluorescence data above 740nm with a 740nm long-pass filter (Schumacher et al., 2014; Trinh le et al., 2007). Transmitted images of chromogenic substrate are shown in gray-scale images (*Spp1*, Figure S6A) and fluorescence of the color development reagent (BM purple) is shown as cyan (*Spp1*, Figures 5C, S6A) or magenta (*Myh9*, Figure 5E).

For the RNAscope single transcript quantification, Z-stacks of ~6 *µ*m from multiple (9-12 per sample) focal planes were obtained on 60x water immersion (WI) objective at Nyquist resolution on the Nikon Ti-E A1R HD confocal with the resonant scanner. The images were processed with NIS-Elements AR 5.2.00 artificial intelligence denoise algorithm (https://www.microscope.healthcare.nikon.com/products/confocal-microscopes/a1hd25-a1rhd25/nis-elements-ai) and stitched into the single image with NIS-Elements AR stitching tool. All images within an experimental group were obtained with the same optical configuration. We first identified individual renal tubules based on the marker gene expression using the “Manual Surfaces” algorithm in Bitplane Imaris 9.3.1. (Chhipa et al., 2018). Then, we identified the transcripts using the “Spots” algorithm, with identical spot diameter and quality used for the experimental groups and the controls. To quantify the transcript number per tubule, we used the Matlab XT function “split spots into surfaces”. We normalized the obtained transcript number to the tubule volume and presented the average of all renal tubules (~50-70 tubules) captured in the image.

### Statistical Analysis and Reproducibility

The gene expression changes identified by the scRNAseq were reproduced in the separate cohort of experimental mice of the same strain, age and sex, treated with the identical UIR procedure. Three to six mice per group were used for the qPCR and Western blot analysis. Data were presented as Mean ± SEM. To determine the statistical significance, P values were generated using one-way ANOVA with Bonferroni and Holm test with * p<0.05 representing the statistically significant difference. The significance is shown compared to the control group. For the RNAscope, 9-12 Z-stack images per animal were analyzed (n=1 animal per time point). Transcript numbers were quantified using Bitplane Imaris 9.3.1. as described above; data were presented as Mean ± SEM and analyzed with Student’s *t*-test or one-way ANOVA with Bonferroni and Holm test when comparing the two or multiple conditions, respectively; p<0.05 represented the statistically significant difference. For the correlation analysis, R squared was obtained and analyzed with Pearson correlation test; p<0.05 represented the statistical significance. The gene ontology (GO) analysis of biological processes was performed in the ToppGene Suite (Chen et al., 2009) with 0.05 p value cutoff. The gene clusters enriched in renal cell populations were generated with the ToppCluster (Kaimal et al., 2010) with Bonferroni correction and 0.05 p value cutoff using the gene level Fruchterman-Reingold algorithm.

### Data and Code Availability

The scRNAseq data were deposited at GEO accession number: GEO: GSE139506.

## Supplemental Information

Figures S1-S6.

**Table S1.** UIR Day1 vs Control gene expression analysis. Related to Figures 1, 2, 4.

**Table S2.** Dedifferenriated Prox UIR Day 7 and 11 Marker Gene. Related to Figure 3.

**Table S3.** "Injured Prox" and "Mixed Identity Cells" Differential Analysis. Related to Figure 3.

**Table S4.** Receptor-Ligand Interactions in Control and UIR Day 4 Kidney. Related to Figure 6.

**Table S5.** qPCR primers, riboprobes, CISH probe primer sequences. Related to Methods.

**Movie 1.** RNAscope Control proximal tubule, *Vim* (green), *Col18a1* (red), *Slc34a1* (purple), 60x Nyquist zoom 0.07 *µ*m/px, scale 20 *µ*m. Related to Figure 6.

**Movie 2.** RNAscope UIR Day 1 proximal tubule, *Vim* (green), *Col18a1* (red), *Slc34a1* (purple), 60x Nyquist zoom 0.07 *µ*m/px, scale 20 *µ*m. Related to Figure 6. Note elevated *Vim* and *Col18a1* in the *Slc34a1* positive tubule.

