## Supplementary material for "Single Cell Profiling of Acute Kidney Injury Reveals Novel Transcriptional Signatures, Mixed Identities and Epithelial-to-Stromal Crosstalk": Figures S1-S6

#### Supplemental Figure Legends

Figure S1. Related to Figure 1.
(A) Feature Plots for renal cell populations identified in the Control kidney by the scRNA-seq. Both injury markers *Havcr1* and *Lcn2* are nearly absent.
(B) Feature Plots identify UIR Day 1 cell populations, including sub-clusters of injured (high *Kim1, a.k.a. Havcr1,* and low *Slc34a1*) and cycling (*Mki67* positive) proximal tubules, the injured *Lcn2*-positive distal tubule, loop of Henle and collecting duct. The “Mixed Identity Cells” are positioned between the proximal tubules, distal tubules, loop of Henle and collecting duct.
(C) RNAscope showing *Lcn2* elevation in the distal nephron tubule and collecting duct of UIR Day 1. *Lcn2* (green), *Slc34a1* (purple), 4x (2500um scale) and 10x (500um scale).
(D) H&E staining reveals AKI induced tubular dilation (blue pointers) and cast formation (yellow pointers) not detectable in the Control. 40x, 100um scale.
(E) GO Biological Process of “Injured Prox” UIR Day 1 vs Control, –log2(pValue).

Figure S2. Related to Figures 1, 2.
(A) Feature plots show the expression of the loop of Henle marker *Umod* and the collecting duct marker *Aqp2* in the UIR Day 1. The intermediately located “Mixed Identity Cells” pointed with the arrow show overlapping expression of both markers.
(B) Combined feature plot shows the overlapping expression of the injury markers Kim1 and Lcn2 in the UIR Day 1 “Mixed Identity Cells”.
(C) Heatmap shows the relative marker gene expression in the Control renal cell populations.

Figure S3. Related to Figure 3.
(A) H&E shows the pronounced renal tubular injury at Day 2, which starts resolving at Day 4 and 7. UIR Day 11 and 14 exhibit normal renal histology. 40x. Tubular dilation (blue pointers), cast formation (yellow pointers).
(B) qPCR shows *Kim1* and *Lcn2* expression over the AKI course, Mean ± SEM, one way ANOVA with Bonferroni, ** pValue<0.01 compared to Control.
(C) Western blots image and quantification show Kim1 and Lcn2 expression over the AKI course, Mean ± SEM, one way ANOVA with Bonferroni, ** pValue<0.01 compared to Control.
(D) UMAP shows renal cell populations in the UIR Day 11.
(E) *Slc34a1* CISH, UIR Day 11, 4x, 40x.
(F) *Kim1* (green), *Aqp2* (red), *Slc34a1* (purple) RNAscope, UIR Day 11, 4x, 60x.
(G) GO Biological Process analysis of the UIR Day 14 marker genes, -log(2) pValue.
(H) GO Biological processes of genes overlapping in the “Mixed Identity Cells” at Day 1, 2 and 4, -log(2) pValue. Renal developmental pathways highlighted in salmon color.
(I) ToppCluster analysis of the 99 genes overlapping between the UIR Day 1, 2 and 4 in the “Mixed Identity Cells” shows the enrichment of kidney and epithelium development biological processes. *Sox4* and *Cd24a* are highlighted in red. The analysis is done with 0.05 pValue cutoff and Bonferroni correction using the Fruchterman-Reingold algorithm.
Scale 4x, 2500um, 40x, 100um, 60x, 25um.

Figure S4. Related to Figure 4.

(A) RNAscope with *Slc34a1* (pink), *Sox4* (green) and *Kim1* (white) probes, DAPI (blue), UIR Day 1, 2, 7. 60x Nyquist zoom, 0.14 µm/px, Maximal Intensity Projection (MaxIP) from Z-stack, scale 50 µm.
(B) IMARIS quantification of *Sox4* in the UIR Day 1, 2, 4 and 7 vs Control *Slc34a1* vs *Kim1*-positive renal tubules, n=12 Z-stacks (50-70 tubules) per group, Mean ± SEM, Student's *t* test, **** p<0.0001, n.s. – not significant.

Figure S5. Related to Figure 4.
(A) 60x RNAscope images show *Lcn2* (green) and *Cd24a* (pink) colocalization at UIR Day 2, 4 and 7 highlighted with pointers, 60x Nyquist zoom, 0.21 µm/px, Maximal Intensity Projection (MaxIP) from Z-stack, scale 50µm.
(B) IMARIS quantification of *Cd24a* in the UIR Day 1, 2, 4 and 7 vs Control *Slc34a1*-positive renal tubules, n=9 Z-stacks (~60 tubules) per group, Mean ± SEM, analyzed with one way ANOVA with Bonferroni and Holm, ** p<0.01 compared to the Control.
(C) IMARIS quantification of *Cd24a* in the UIR Day 1, 2, 4 and 7 *Lcn2* vs Control *Slc34a1*-positive renal tubules, n=9 Z-stacks (~60 tubules) per group, Mean ± SEM, Student's *t* test, **** p<0.0001.
(D) Pearson's correlation analysis of *Cd24a* and *Lcn2* transcript numbers in UIR Day 1, n=9 Z-stacks (~60 tubules).
(E) Cd24a immunohistochemistry at UIR Day 1 vs Control.

Figure S6. Related to Figure 5.
(A) *Spp1* CISH (cyan) co-stained with Krt8 IF (red) at UIR Day 1, 4 and 14 vs Control. Transmitted detector (TD) shows widefield *Spp1* CISH signal. 20x, 200um scale.
(B) Feature Plots show *Vim* and *Col18a1* in the UIR Day 4 vs Control renal cell populations.
(C and D) qPCR shows Vim and Col18a1 gene expression changes over the AKI course, normalized fold change presented as Mean + SEM, analyzed with one way ANOVA with Bonferroni and Holm, * p<0.05, ** p<0.01 compared to the Control.
(E) RNAscope images show strong *Slc34a1* (purple) proximal tubule marker and low *Vim* (green) and *Col18a1* (red) expression in the interstitial and periglomerular spaces of the Control kidney (white pointers); UIR causes significant *Slc34a1* decline and *Vim* and *Col18a1* elevation in the proximal tubules and in the interstitium, which resolves at Day 11 and 14, with some remaining expression around the glomeruli. 10x (500um scale).

#### Figure S1

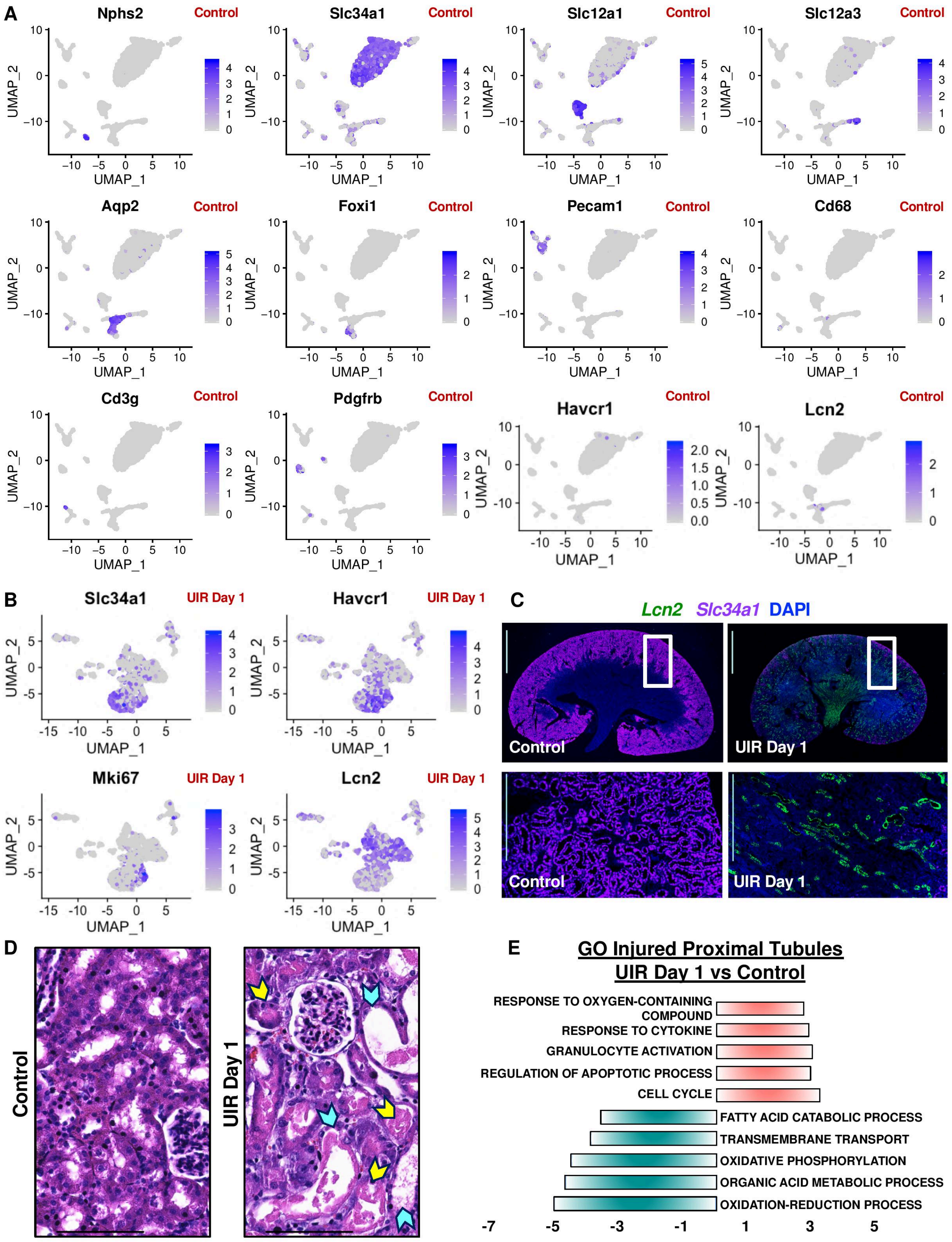
A
Nphs2
Control
Slc34a1
Slc12a1
Slc12a3
Aqp2
Foxi1
Pecam1
Cd68
Cd3g
Pdgfrb
Havcr1
Lcn2
UMAP_1
UMAP_2
B
UIR Day 1
Mki67
C
Lcn2 Slc34a1 DAPI
Control
UIR Day 1
D
E
GO Injured Proximal Tubules
UIR Day 1 vs Control
RESPONSE TO OXYGEN-CONTAINING COMPOUND
RESPONSE TO CYTOKINE
GRANULOCYTE ACTIVATION
REGULATION OF APOPTOTIC PROCESS
CELL CYCLE
FATTY ACID CATABOLIC PROCESS
TRANSMEMBRANE TRANSPORT
OXIDATIVE PHOSPHORYLATION
ORGANIC ACID METABOLIC PROCESS
OXIDATION-REDUCTION PROCESS
-7
-5
-3
-1
1
3
5

### Figure S2

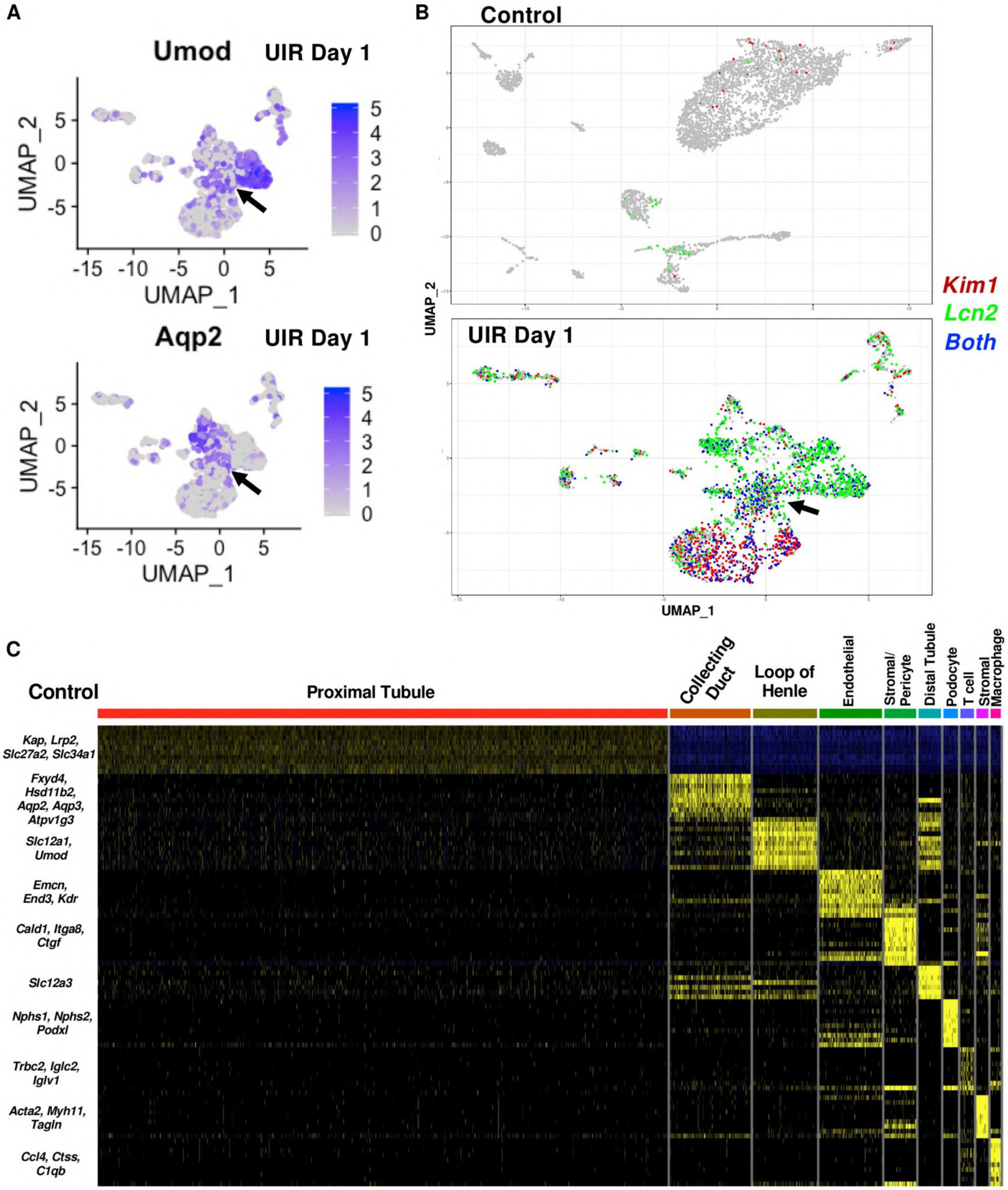
A
Umod UIR Day 1
Aqp2 UIR Day 1
UMAP_1
UMAP_2
B
Control
UIR Day 1
Kim1
Lcn2
Both
UMAP_1
UMAP_2
C
Control
Proximal Tubule
Collecting Duct
Loop of Henle
Endothelial
Stromal/ Pericyte
Distal Tubule
Podocyte
T cell
Stromal
Macrophage
Kap, Lrp2, Slc27a2, Slc34a1
Fxyd4, Hsd11b2, Aqp2, Aqp3, Atpv1g3
Slc12a1, Umod
Emcn, End3, Kdr
Cald1, Itga8, Ctgf
Slc12a3
Nphs1, Nphs2, Podxl
Trbc2, Iglc2, Iglv1
Acta2, Myh11, Tagln
Ccl4, Ctss, C1qb

### Figure S3

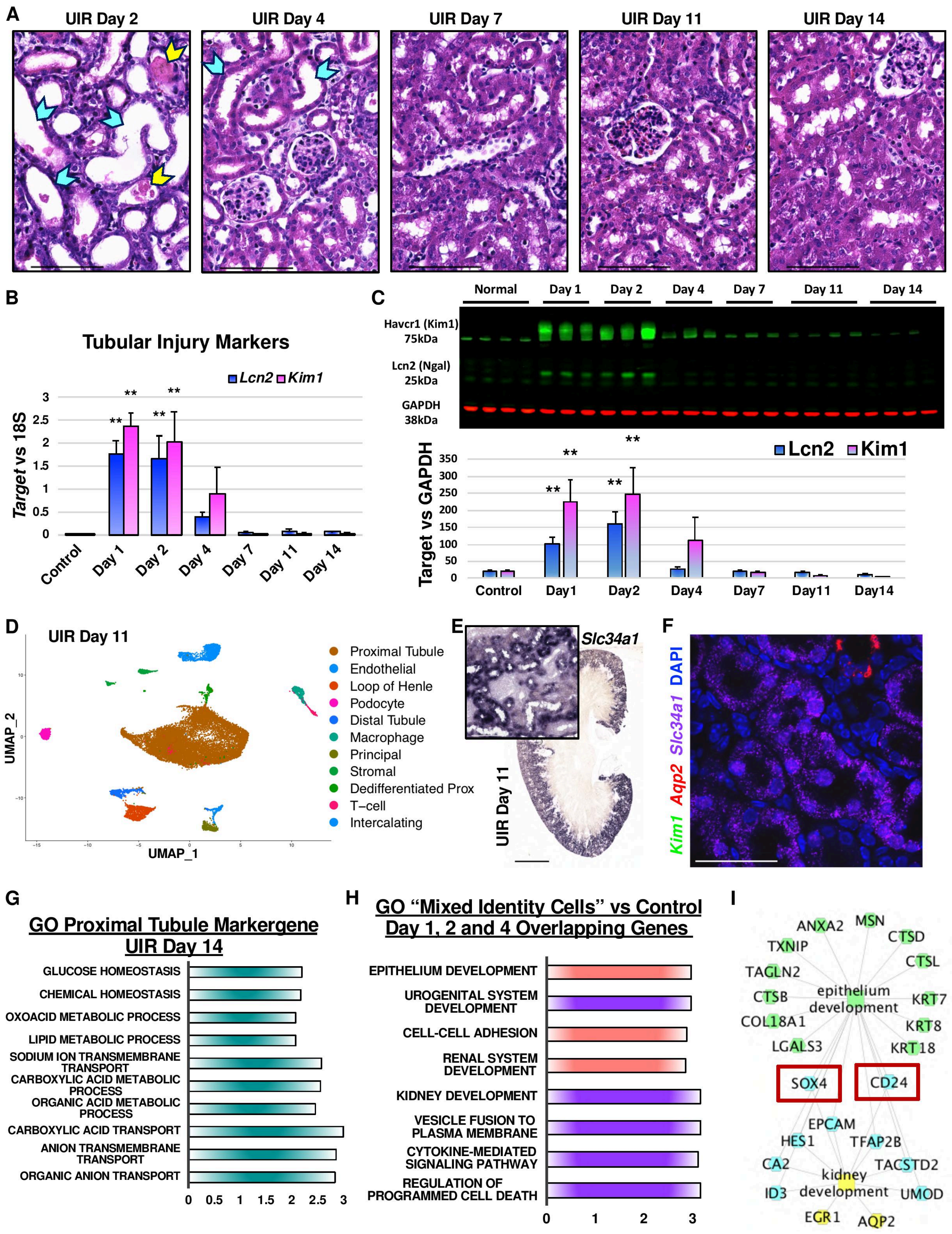

A
UIR Day 2
UIR Day 4
UIR Day 7
UIR Day 11
UIR Day 14
B
Tubular Injury Markers
Lcn2
Kim1
Target vs 18S
Control
Day 1
Day 2
Day 4
Day 7
Day 11
Day 14
C
Normal
Day 1
Day 2
Day 4
Day 7
Day 11
Day 14
Havcr1 (Kim1) 75kDa
Lcn2 (Ngal) 25kDa
GAPDH 38kDa
Lcn2
Kim1
Target vs GAPDH
Control
Day1
Day2
Day4
Day7
Day11
Day14
D
UIR Day 11
UMAP_2
UMAP_1
Proximal Tubule
Endothelial
Loop of Henle
Podocyte
Distal Tubule
Macrophage
Principal
Stromal
Dedifferentiated Prox
T–cell
Intercalating
E
Slc34a1
UIR Day 11
F
Kim1 Aqp2 Slc34a1 DAPI
G
GO Proximal Tubule Markergene UIR Day 14
GLUCOSE HOMEOSTASIS
CHEMICAL HOMEOSTASIS
OXOACID METABOLIC PROCESS
LIPID METABOLIC PROCESS
SODIUM ION TRANSMEMBRANE TRANSPORT
CARBOXYLIC ACID METABOLIC PROCESS
ORGANIC ACID METABOLIC PROCESS
CARBOXYLIC ACID TRANSPORT
ANION TRANSMEMBRANE TRANSPORT
ORGANIC ANION TRANSPORT
H
GO “Mixed Identity Cells” vs Control Day 1, 2 and 4 Overlapping Genes
EPITHELIUM DEVELOPMENT
UROGENITAL SYSTEM DEVELOPMENT
CELL-CELL ADHESION
RENAL SYSTEM DEVELOPMENT
KIDNEY DEVELOPMENT
VESICLE FUSION TO PLASMA MEMBRANE
CYTOKINE-MEDIATED SIGNALING PATHWAY
REGULATION OF PROGRAMMED CELL DEATH
I
ANXA2
MSN
CTSD
TXNIP
CTSL
TAGLN2
CTSB
epithelium development
KRT7
COL18A1
KRT8
LGALS3
KRT18
SOX4
CD24
EPCAM
HES1
TFAP2B
CA2
TACSTD2
kidney development
ID3
UMOD
EGR1
AQP2

#### Figure S4

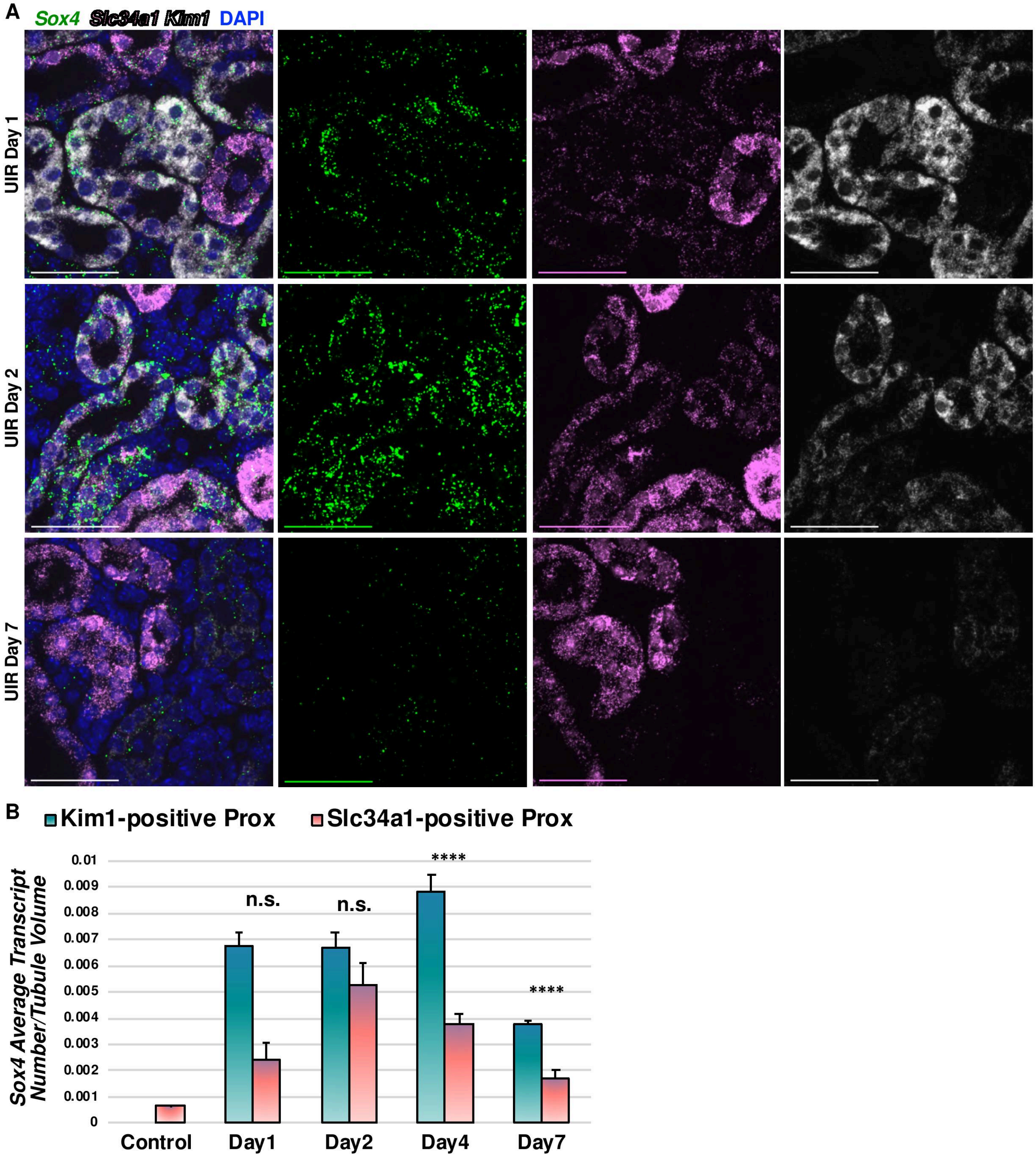

A
Sox4 Slc34a1 Kim1 DAPI
UIR Day 1
UIR Day 2
UIR Day 7
B
Kim1-positive Prox
Slc34a1-positive Prox
Sox4 Average Transcript Number/Tubule Volume
0.01
0.009
0.008
0.007
0.006
0.005
0.004
0.003
0.002
0.001
0
n.s.
n.s.
****
****
Control
Day1
Day2
Day4
Day7

#### Figure S5

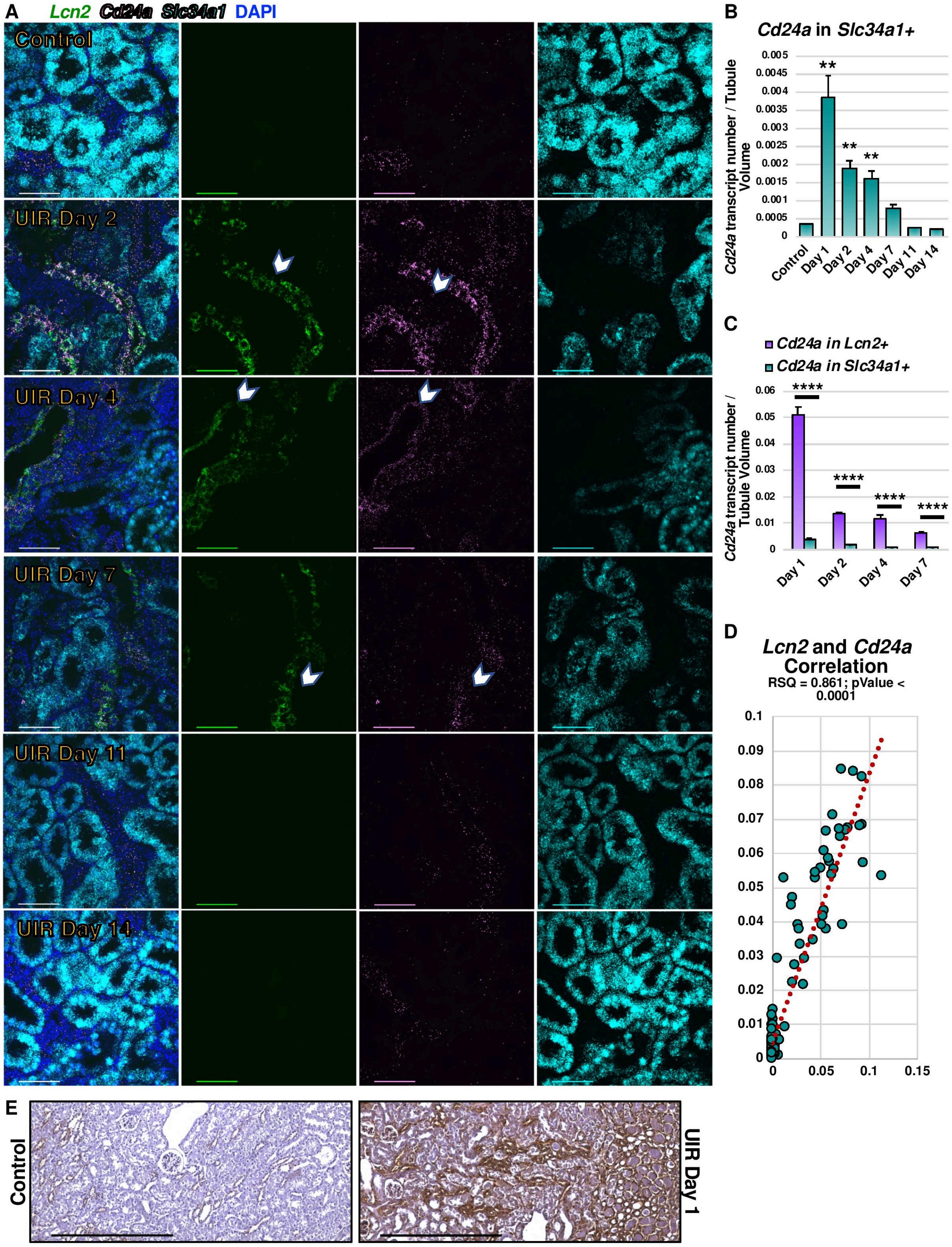
A
Lcn2 Cd24a Slc34a1 DAPI
Control
UIR Day 2
UIR Day 4
UIR Day 7
UIR Day 11
UIR Day 14
B
Cd24a in Slc34a1+
Cd24a transcript number / Tubule Volume
Control
Day 1
Day 2
Day 4
Day 7
Day 11
Day 14
C
Cd24a in Lcn2+
Cd24a in Slc34a1+
Cd24a transcript number / Tubule Volume
D
Lcn2 and Cd24a Correlation
RSQ = 0.861; pValue < 0.0001
E
Control
UIR Day 1

#### Figure S6

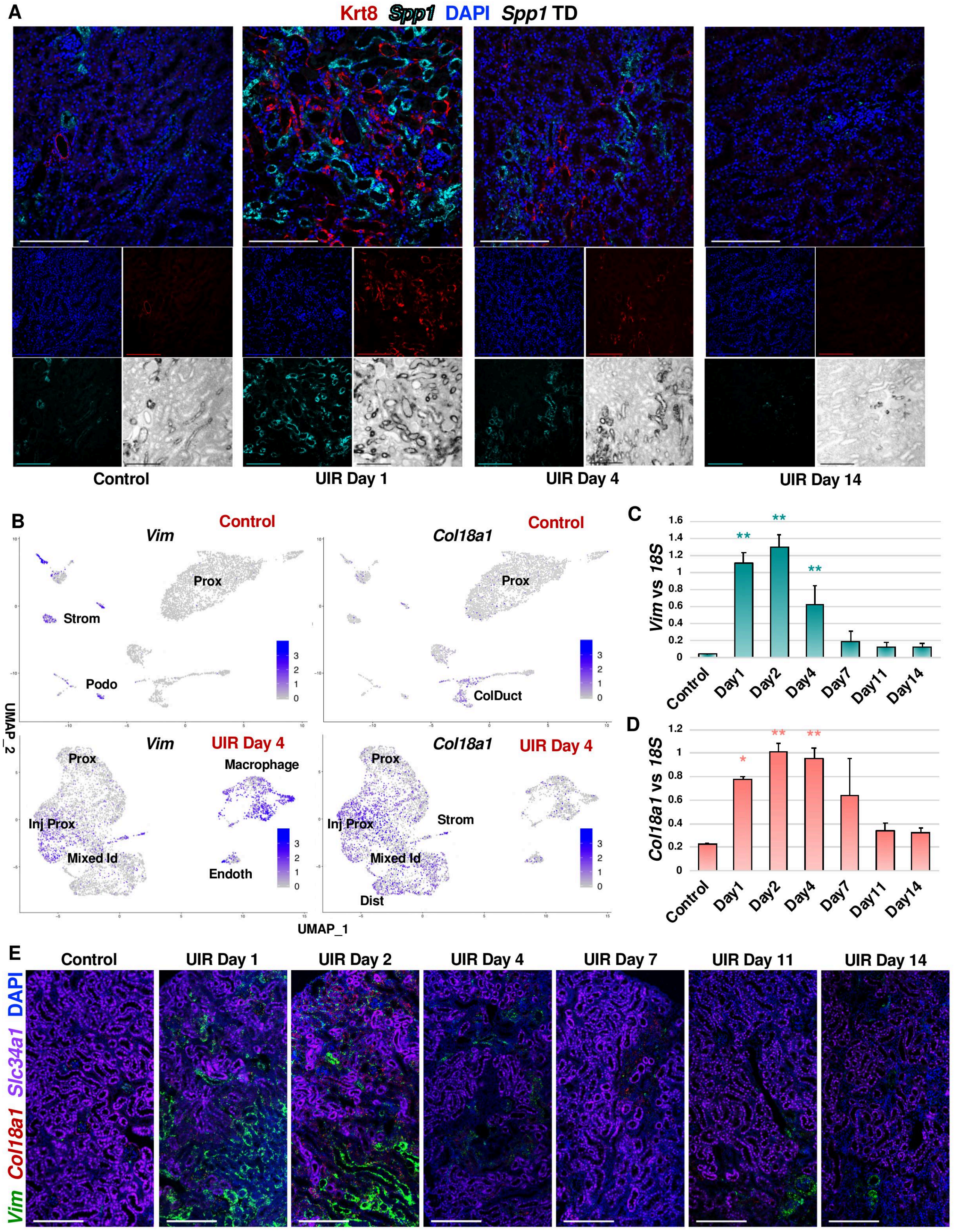

A
Krt8 Spp1 DAPI Spp1 TD
Control
UIR Day 1
UIR Day 4
UIR Day 14
B
Vim
Control
Prox
Strom
Podo
Col18a1
Control
Prox
ColDuct
UMAP_2
Vim
UIR Day 4
Prox
Macrophage
Inj Prox
Mixed Id
Endoth
Col18a1
UIR Day 4
Prox
Inj Prox
Strom
Mixed Id
Dist
UMAP_1
C
Vim vs 18S
Control
Day1
Day2
Day4
Day7
Day11
Day14
D
Col18a1 vs 18S
Control
Day1
Day2
Day4
Day7
Day11
Day14
E
Vim Col18a1 Slc34a1 DAPI
Control
UIR Day 1
UIR Day 2
UIR Day 4
UIR Day 7
UIR Day 11
UIR Day 14
